# *Mycobacterium tuberculosis* Genes Needed for Intra-alveolar Resuscitation After Airborne Transmission

**DOI:** 10.1101/2025.09.15.676370

**Authors:** Prabhat Ranjan Singh, Martin Gengenbacher, Saurabh Mishra, Adrian Jinich, Xiuju Jiang, Felipe Tsang, Melissa Cristaldo, Michael DeJesus, Kyu Y. Rhee, Robert J. Kaner, Philip L. Leopold, Ronald G Crystal, Dipak Kathayat, Brian C. VanderVen, Carl Nathan

## Abstract

*Mycobacterium tuberculosis* (Mtb) must survive multiple changes in environment for aerosol transmission. Our genome-wide screen for rehydration of modeled aerosols in surrogate pulmonary alveolar lining fluid identified a survival-sustaining role for 22 genes not required for survival during earlier stages of transmission, 20 of which are non-essential in routine culture. Thirteen genes were also needed for Mtb to survive in alveolar macrophages after passing through the earlier stages of transmission. Nine 9 of the 13 were needed for full infectivity of aerosols in mice. Seven of the genes sustaining the intra-alveolar survival of aerosolized Mtb are likely to regulate Mtb’s uptake, catabolism or synthesis of lipids as regulated by cAMP. Our results reveal a dynamic form of conditional essentiality that only emerges after bacteria experience a sequence of other pathophysiologically relevant conditions. These findings enlarge Mtb’s candidate transmission survival genome with stage-specific genes encoding potential targets for blocking tuberculosis transmission.

## Introduction

A quarter century after publication of the genome sequence of Mtb ^1^, the world’s leading cause of death from infection ^2^, the function of about half of the pathogen’s ∼4100 genes remains unknown and an implausibly large proportion. Moreover, ∼74%, are considered non-essential ^3–5^. We reasoned that the knowledge gap may reflect the inability of conventional culture systems and mouse models to present Mtb with certain conditions that have shaped its evolution. Among these is likely to be aerosol transmission ^6–9^ , a multi-stage process during which the bacilli encounter strikingly different liquid and/or gaseous environments ^8,9^.

To our knowledge, there is as yet no way to conduct a genome-wide screen of Mtb genes during aerosol transmission in animals. We instead sought to identify Mtb’s candidate transmission survival genome using an in vitro model, while acknowledging the histopathologic diversity of infected tissues from which Mtb may exit from a human host, the complexity and variety of the liquid and gas phases in those sites, differences in air temperature, relative humidity, sunlight irradiance and inter-individual distances during the flight of infectious aerosols, and the complexity of the pulmonary alveolar lining fluid in which inhaled Mtb aerosols initiate infection. We reasoned that a screen under sequential pathophysiologically relevant conditions could nominate candidate genes, some of which could then be queried individually in animal models.

The first round of this effort required the development of a bicarbonate-buffered model aerosol fluid (MAF) based on the composition of caseum from necrotic TB lesions as it might be fluidized by bronchial secretions to the viscosity required to form aerosols that are small enough (< ∼ 5 μm) to remain suspended for prolonged periods, travel more than about a meter and reach pulmonary alveoli in new hosts. Using MAF and a sequence of atmospheres, we modeled 3 stages of TB transmission^9^. As schematized in **Fig. 1**, stage 0 refers to conventional log phase growth in 7H9 bacteriologic medium under 21% O_2_ and 5% CO_2_; stage 1 models residence for 2 weeks in a hypoxic (0.2% O_2_), eucapnic (5% CO_2_), necrotic pulmonary lesion; stage 2 models extrusion of lesional contents into an airway for 2 weeks (10% O_2_, 5% CO_2_); and stage 3 represents exposure of droplets to air outside the host (21% O_2_, 0.4% CO_2_), where Mtb experiences desiccation, cooling, oxidation, hypocapnia and alkalinization, against which MAF affords substantial protection ^9,10^. We subjected a genome-wide CRISPRi library of Mtb to all 3 stages in sequence. This led to the nomination of hundreds of genes as members of Mtb’s candidate transmission survival genome ^9^.

**Figure 1.**
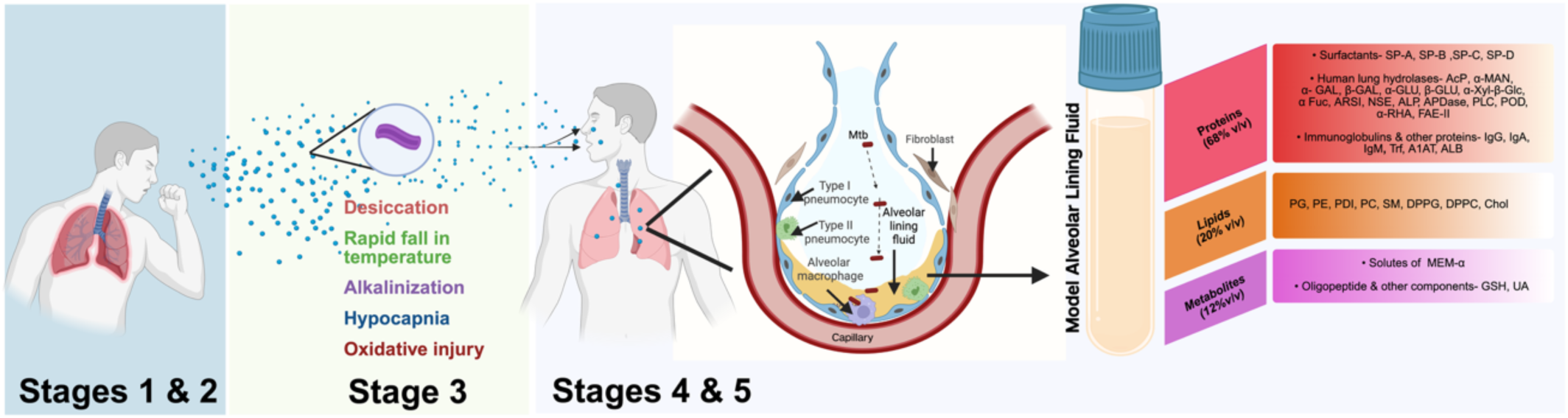
Sequential stages in the transmission of Mtb. The schematic illustrates the progression of Mtb from its residence in a closed, hypoxic pulmonary cavity (stage 1) to its release into the airway (stage 2) and subsequent aerosolization (Stage 3). During the airborne phase, the bacilli experience desiccation, cooling, alkalinization and oxidative stress before inhalation by a new host. Upon reaching the pulmonary alveolus, Mtb interacts with alveolar lining fluid (stage 4) and is then taken up by alveolar macrophages (stage 5). Model alveolar lining fluid (MALF) mimics human alveolar lining fluid, with proteins, lipids and metabolites at the concentrations given in **Table S1.**

In the present study we extended the sequential stages of modeled transmission by two more stages—rehydration in alveolar lining fluid (ALF), followed by uptake in alveolar macropahges. In stage 4, we passaged the genome-wide CRISPRi library through to the stage of desiccated aerosols and then subjected it to rehydration in the antibacterial milieu of modeled human alveolar lining fluid. We identified 35 genes as required in stage 4, 31 of which are non-essential in standard culture, 21 of which were also not implicated in stages 1-3, and 7 of which have unknown functions. We then queried the 35 for their role in helping Mtb survive engulfment by intra-alveolar macrophages at an air-liquid interface on pulmonary epithelium (stage 5) (**Fig. 1**). Of the 13 genes additionally required in stage 5, we confirmed that 9 support Mtb’s ability to infect mice by inhalation of aerosols of respirable size.

## Results

### Formulation and characterization of MALF

To mimic human ALF, we developed modeled alveolar lining fluid (MALF), not to be confused with MAF, which has an extensively different composition ^9^. We inferred the composition of ALF from multiple reports on the composition of bronchoalveolar lavage fluid (BALF) from healthy donors ^11–29^. Few of the literature reports identified the same components and we included all components that these reports identified. Therefore, MALF is considerably more complex than other simulations of ALF. The verisimilitude of ALF simulations is limited by inter-donor variability ^30,31^, by the unknown degree to which lavage with physiologic saline recovers lipids, and by the need to correct for the dilution of analytes by bronchoalveolar lavage ^32^. **Fig. 1** lists the lipids, proteins and metabolites comprising MALF; Supplemental Methods and **table S1** give their concentrations. In contrast to standard Mtb culture media, the osmolality of MALF (∼330 mOsm/kg) was close to that of human body fluids (275 to 295 mOsm/kg) and mammalian cell culture fluids (**fig. S1**). The pH of MALF was ∼7.4.

To test whether MALF is a functionally relevant mimic of human ALF, we compared it to BALF collected at bronchoscopy from three healthy, non-smoking adults **(table S2)**. We estimated the fold dilution of ALF by the saline used for lavage by assuming equilibration of urea between serum and ALF ^32^ (**fig. S2**). With relative urea concentrations as a guide, we reduced the volume of BALF by filtration through a 3 kDa Centricon filter at 4 °C by an amount sufficient to reverse the dilution of macromolecules. In another departure from previous efforts to model ALF, we restored many of the analytes lost by filtration by dialyzing the filtrate overnight against MEM-α, a mammalian cell culture medium (**Fig. 2A**). We termed the dialysand “reconstituted BALF” (rcBALF). The pH of rcBALF from 3 healthy adult donors was ∼7.5 and their osmolality was ∼300 mOsm/kg (**fig. S1**), comparable to that of MALF. MALF was not cytotoxic to the donor cells recovered by centrifugation of BALF (**fig. S3**).

**Figure 2.**
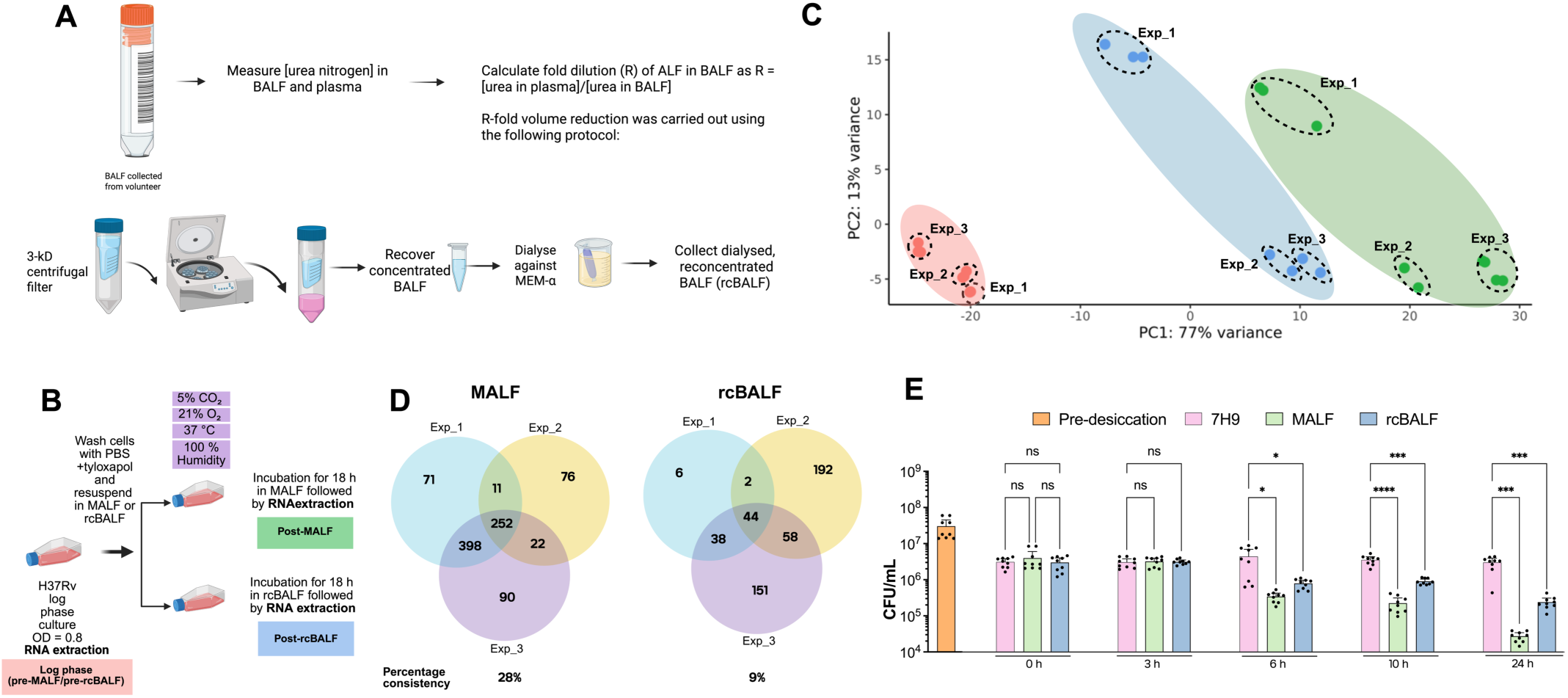
Characterization of MALF with reference to BALF. **(A)** Workflow for producing re-concentrated BALF (rcBALF) to correct for the dilution of ALF by lavage of lung segments with 0.9% saline. **(B)** Schematic of the RNA-seq experiment using log-phase Mtb cultures. **(C)** PCA plot comparing transcriptomes of log-phase Mtb in conventional culture medium (7H9) with Mtb incubated for 18 h in MALF or rcBALF. **(D)** Statistical comparison of RNA-seq results from three independent experiments each in MALF and in rcBALF, using rcBALF prepared from 3 different donors. (padj value ≤ 0.05, log_2_ FC ≥ 1.5 or log_2_ FC ≤ -1.5).The central intersect in each Venn diagram includes all genes whose expression changed in either direction in all 3 experiments. includes all Percentages below the Venn diagrams indicate the proportion of the differentially transcribed genes for which the changes were consistent in each experiment. **(F)** Survival of Mtb that had passed through stages 1-3 before incubation in 7H9, rcBALF or MALF. Data are means ± SEM from 3 independent experiments.

Using RNA-seq, we compared the transcriptome of Mtb in MALF to the transcriptome of Mtb in rcBALF. In a first set of experiments, we used Mtb that had passed through the first three stages of modeled transmission, the first two stages of which induce massive transcriptional adaptations ^9^. The transcriptomes of Mtb progressing out of stages 1, 2, and 3 into MALF or into rcBALF **(fig. S4A)** were almost indistinguishable from each other **(fig. S4B**) and from the transcriptome of Mtb harvested directly from stage 3 (desiccation) into 2.5 M guanidinium thiocyanate or into MEM-α (f**ig. S4C**). The lack of consistently differentially expressed genes going from stage 3 (desiccation) to stage 4 (rehydration) in either medium under these conditions suggested functional similarity between MALF and rcBALF.

Reasoning that a larger number of differentially expressed genes could allow a more stringent comparison of Mtb’s responses to MALF and rcBALF, we repeated these studies but omitted passage of conventionally cultured Mtb through stages 1-3 before resuspending the bacteria for 18 hours in MLF or rcBALF under 21% O_2_, 5% CO_2_ (**Fig. 2B**). In three independent experiments, 109 genes were consistently upregulated and 110 were consistently downregulated in MALF-exposed Mtb, compared to 44 genes consistently upregulated and none consistently downregulated in rcBALF-exposed Mtb (**Fig. 2C**). Notably, 41 of the 44 genes differentially expressed in rcBALF (93%) were also consistently differentially expressed in MALF-exposed Mtb **(Fig. S5**). There was greater variability in transcriptomic changes of Mtb incubated in rcBALF prepared from 3 donors (9% congruence) than in MALF prepared on 3 occasions (28% congruence) (**Fig. 2D**), although variability was considerable in both cases. For MALF, we speculate that variability may be due to the difficulty in minimizing the oxidation of lipids during preparation. For rcBALF, we speculate that variability may in part be related to differences among the donors, including the quality of the air they habitually breathe, but may also reflect differences in recovery of lipids from ALF by lavage with saline, in which lipids are insoluble.

In sum, once Mtb had passed through the first three stages of modeled transmission, we could detect no substantive transcriptional differences between Mtb in desiccated droplets undergoing rehydration in MALF or in rcBALF. When we used Mtb from conventional culture, MALF induced a broader and more consistent gene expression response than rcBALF, including nearly all of the genes differentially expressed in rcBALF.

When Mtb H37Rv was taken from log phase culture in 7H9 broth and incubated in MALF or rcBALF, there was no detectable loss of viability by 18 h (**fig. S6A**), notwithstanding the antimicrobial properties of human ALF ^29–31,33^. We then repeated the viability test using Mtb that had passed through stages 1-3 of the transmission model, including desiccation, in MAF (not MALF). Rehydration in 7H9 medium and culture under 21% O_2_ with 5% CO_2_ revealed a ∼1-log₁₀ loss in viability of Mtb coming out of the desiccation stage, as seen earlier ^9^, but there was no further reduction in colony-forming units (CFU) over the next 24 hours in 7H9. In contrast, upon rehydration in MALF or rcBALF, Mtb’s viability was maintained for at least 3 hours but fell by the next time point studied, 6 hours (**Fig. 2E**). Although this is consistent with the expected antimicrobial effect of ALF, 3 hours is well past the 10 minutes that were required for alveolar macrophages to phagocytize all the adenovirus particles that reached pulmonary alveoli in mice ^34^. As summarized in **fig. S6B, C**, multiple components of MALF appeared to interact to confer its delayed anti-mycobacterial effect. Beyond 3 hours, desiccation-stressed Mtb showed a similar survival deficit in rcBALF as in MALF (**Fig. 2E**), further supporting that MALF recapitulates some key features of rcBALF.

### Genes sustaining Mtb’s survival during modeled transition from aerosolization to inhalation

Next, working with triplicate samples, we passed the genome-wide CRISPR interference (CRISPRi) library targeting 4,014 genes in Mtb **(fig. S7)** from the previously described stages 1, 2 and 3 ^9^ into stage 4 by resuspending the desiccated droplets of stage 3 in MALF for 18 hours under 21% O₂, 5% CO₂ at 37 °C. We sequenced the sgRNAs in surviving cells in one portion of the samples directly after the 18 h incubation in MALF and in another portion after allowing for biomass expansion through 5 generations of outgrowth in 7H9 medium under 21% O₂, 5% CO₂. We determined the abundance of each sgRNA in the post-MALF samples (stage 4, with and without outgrowth) vs. the stage 0 samples in which dCas9 had been induced for 5, 10 or 20 generations **(fig. S7)**. Using MAGeCK ^35^, we applied thresholds of ≤ –1.5 log_2_ fold change and p ≤ 0.05 to represent a significant difference. We identified 784 genes with a number of significant differences (NOSD) ≥ 1 in these comparisons as candidates for aiding Mtb’s survival in transition to stage 4 (inhalation) (**table S3**).

We identified 35 genes that appeared to sustain Mtb’s survival in stage 4 (resuscitation) but not in stage 3 (desiccation). Of the 35, 21 appeared to play a role uniquely in the 4th stage, while 12 genes also appeared to play a role in stage 1, and two genes also appeared to play a role in stage 2. The 35 genes included 31 conventionally nonessential genes and 4 conventionally essential genes. Among them are genes representing diverse physiological pathways (**Fig. 3A**).

**Figure 3.**
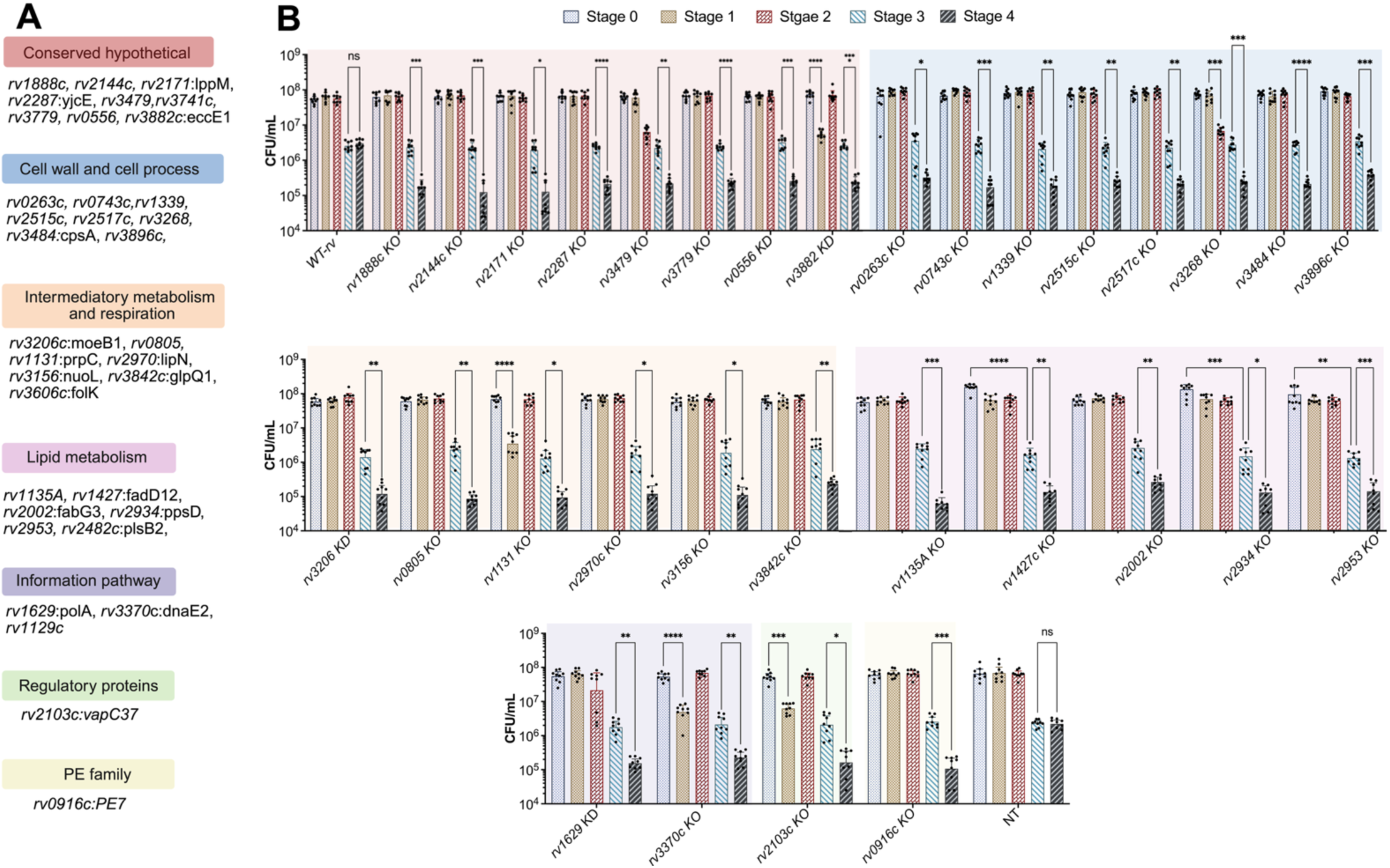
Verification of CRISPRi screening hits using individual knockdown and knockout strains. **(A)** Pathway analysis of the 35 genes chosen for verification as candidates for sustaining Mtb’s survival in the 4^th^ stage (immersion in ALF) but not in the 3^rd^ stage (desiccation of droplets in air) of modeled transmission. **(B)** CFU of control strains—WT (control for knockout strains) and NT (non-targeting control for CRISPRi knockdown (KD) strains)—measured at inoculum (stage 0), after 2 weeks in MAF under 0.2% O₂ (stage 1), followed by 2 weeks in MAF under 10% O₂ (stage 2), after desiccation and recovery in MALF at 0 h post-rehydration (stage 3), and after 18 h incubation in MALF (stage 4). Significance is indicated by asterisks (*p < 0.05, **p < 0.01, ***p < 0.001; ns, non-significant). Results are means ± SEM from 3 independent experiments. Statistical significance was determined by two-way ANOVA followed by Tukey’smultiple comparisons test.

We knocked out the 30 genes of interest that are nonessential under conventional conditions and constructed individual CRISPRi knockdown strains for the 4 essential genes *rv0556, rv1629, rv3206*, and *rv3606*, for which knockouts are nonviable. We were unable to make a knockout of the non-essential gene *rv3882* and so we likewise made a CRISPRi knockdown strain. We confirmed the knockouts by PCR and by whole genome resequencing, which excluded the introduction of adventitious mutations. We tested the survival of the 35 individual mutant strains as they passed sequentially through all 4 stages of modeled transmission (**fig. S8A).** Wild type Mtb and all but 3 of the gene-deficient strains (*rv1131*, *rv3268*, and *rv3882*) showed no survival defect in stages 1 or 2 (**Fig. 3B** & **fig. S8B**). Emerging from stage 3 (desiccation), wild type Mtb and 32 of the knockout and knockdown strains showed ∼1 log₁₀ reduction in CFU compared to stage 0, while the strains with knockout of *rv1427c, rv2171*, or *rv2953* displayed a ∼2 log₁₀ reduction. For 31 of the 35 genes, knockout or knockdown led to a further survival deficit of ∼1 log₁₀ in stage 4 **(Fig. 3B).** The condition-dependent survival defect of the knockout strains was reversed through complementation with the wild type allele, except for the knockouts of *rv1888c*, *rv2171c*, *rv2103* and *rv3268* **(fig. S8B).** Strains deficient in *rv1129c*, *rv2482c, rv3606c*, or *rv3741c* did not show a further survival deficit in stage 4 **(fig. S8C).**

The 31 genes with confirmed phenotypes in stage 4 are candidates for helping aerosolized Mtb resist the antibacterial effects of ALF long enough to take refuge in alveolar macrophages. There, Mtb faces further challenges, as we explored next.

### Role of inhalation-survival genes beyond rehydration in MALF

After Mtb reaches ALF, most of the bacteria are phagocytized, chiefly by pulmonary alveolar macrophages. We next asked which of the genes that were candidates for supporting the survival of Mtb in stage 4 of modeled transmission were additionally required to support Mtb’s survival at the air-liquid interface (ALI) in a culture of alveolar macrophage-like cells atop pulmonary epithelial cells (**Fig. 4A)**. The ALI model was validated by demonstrating the permeability barrier of the epithelial cell monolayer (**fig. S9A)** and its expression of actin and MUC5AC (**fig. S9B**).

**Figure 4.**
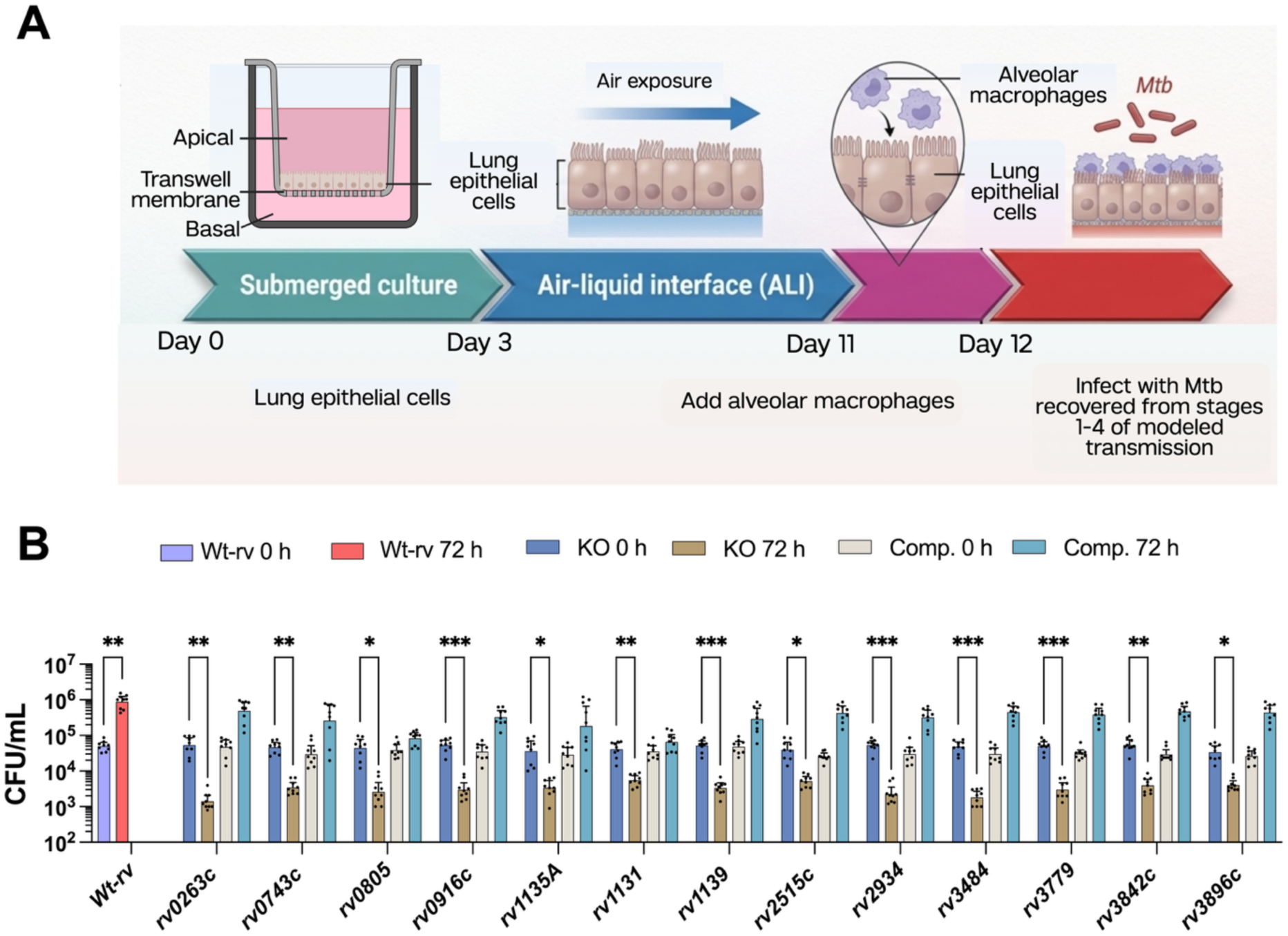
Survival of Mtb wild type and mutant strains in stage 5 of transmission: uptake of Mtb by macrophages at an air-liquid interface (ALI) on pulmonary epithelium. (A) Establishment of the ALI. Lung epithelial cells (Nuli-1) were initially grown in a submerged transwell culture. At day 3, exposure to air initiated the ALI. On day 11, alveolar macrophages (AMJ2-C8) were added atop the epithelial cells. On day 12, the co-culture was infected with Mtb strains recovered from stages 1-4 of modeled transmission. **(B)** Intracellular bacterial growth of wild type H37Rv (Wt-rv) and Mtb strains with the indicated genes knocked out (KO) or complemented (Comp.). CFU were measured at 0 hours and 72 hours post-infection. Significance is indicated by asterisks (*p < 0.05, **p < 0.01, ***p < 0.001; ns, non-significant). Results are means ± SEM from 3 independent experiments. Statistical significance was determined by two-way ANOVA followed by Tukey’smultiple comparisons test.

We passed all 35 knockout or knockdown strains individually through the 4 stages of modeled transmission (**fig. S10A**) and then infected the ALI cultures with 1 Mtb per alveolar macrophage-like cell (stage 5) (**Fig. 4A)**. Over the next 72 hours, 24 mutants showed little or no change in CFU, while the CFU of wild-type Mtb increased 10- fold (**fig. S10B**). In contrast, Mtb strains deficient *in rv0263c, rv0743c, rv0805, rv0916c, rv1131, rv1135A, rv1339, rv2515c, rv2934, rv3484, rv3779, rv3842c* and *rv3896c* each showed ∼ 10-fold reduction in CFU compared to the inoculum, representing a ∼100-fold disadvantage relative to wild-type Mtb at 72 hours (**Fig. 4B**). Complementation of the knockouts with the wild type allele fully reverted the viability defect in 11 of the mutant strains and partially in the *rv0805* and *rv1131* mutant strains.

When the wild type strain, the knockouts of *rv0805* or *rv1339* and their respective complemented strains were taken directly from stage 0 (log phase culture in 7H9 medium under 21% O_2_, 5% CO_2_) and used to infect the alveolar macrophage-like cells in plastic plates, there was no defect in their intracellular replication (**fig. S11A**). When the 13 knock-out strains with phenotypes in alveolar macrophage-like cells at the air-liquid interface were passed through stages 1-3 of modeled transmission and then used to infect human monocyte-derived macrophages at an air-liquid interface ^36^, the phenotype seen with alveolar macrophage-like cells was recapitulated (**fig. S11B**). Thus, the intra-macrophage phenotype associated with these genes was less dependent on the type of macrophages than on the sequence of conditions experienced by the bacteria.

These results suggest that 13 genes may be selectively and critically important for aerosolized Mtb to infect a human by inhalation. Each of them had been considered nonessential, and there are few if any published clues to the function of 6 of them (*rv0263c, rv0743c, rv0916c, rv1135A, rv2515c* and *rv3896c*).

### Infectivity of aerosolized Mtb mutant strains in mice

To explore the relevance of the in vitro model to authentic aerosol transmission, we used a transmission stimulation system (TSS) designed to produce aerosols of respirable dimensions at forces of human exhalation and confirmed that 90% of the aerosol particles it generates from MAF are ≤ 3.3 μm (**Fig. 5A**) ^37^. We passed 13 Mtb knockout strains through stages 1 and 2 of modeled transmission for 2 weeks at each stage, and used these cultures to inoculate 8-10 mice per Mtb strain via aerosols generated in the TSS **(Fig. 5A).** At 24 h post-infection, 9 of the 13 mutant strains (knockouts of *rv0263, rv0805, rv0916, rv1135A, rv1139, rv3484, rv3842, rv3779* and *rv3896*) showed a statistically significant lower pathogen burden in lungs compared to wild type Mtb, indicating reduced infectivity in mice (**Fig. 5B**). Two gene-complemented mutants tested in the TSS (*rv0805* and *rv1339*) showed fully restored infectivity (**Fig. 5C**).

**Figure 5.**
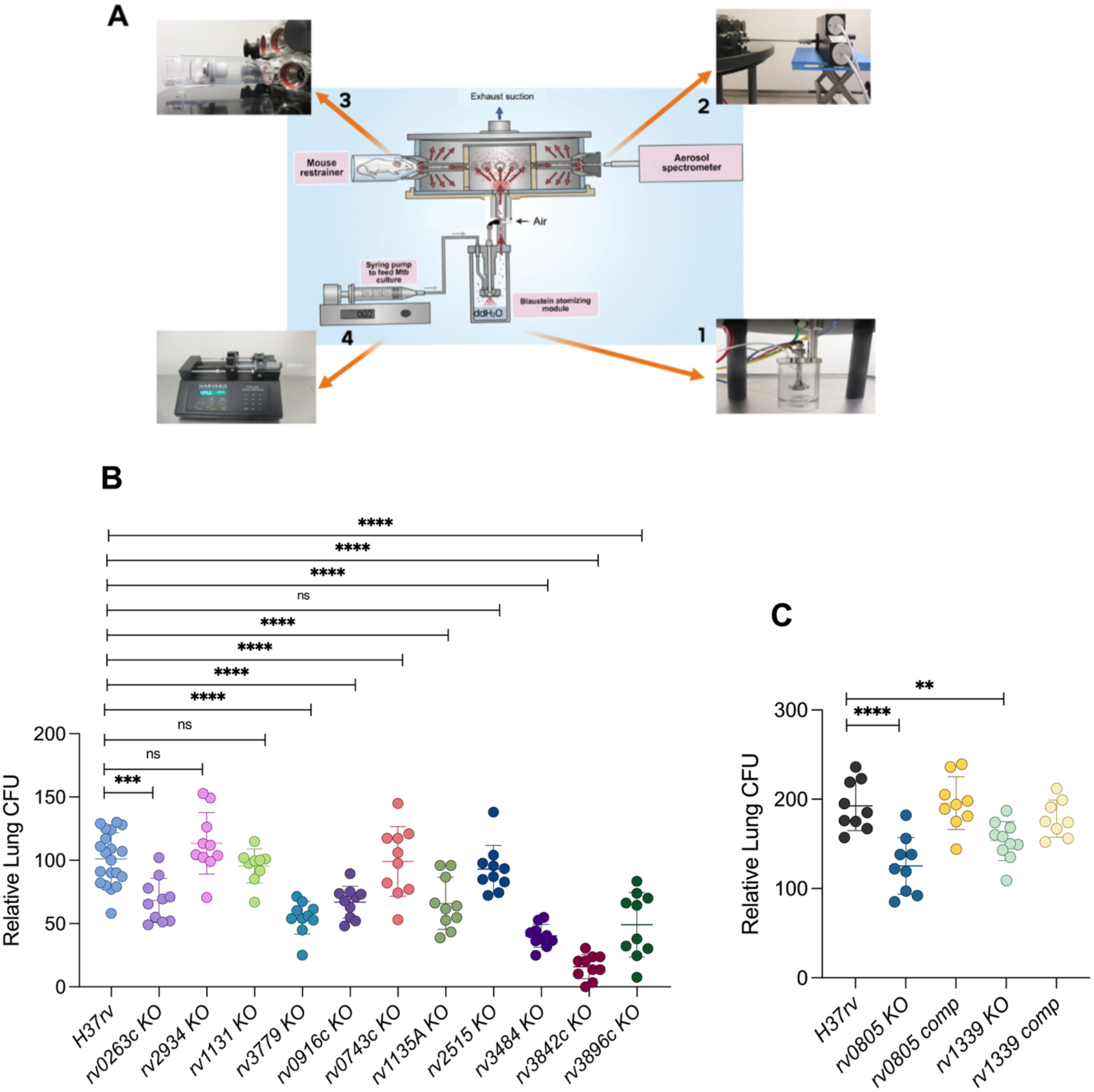
Infectivity of stage 4 candidate transmission survival gene mutants for mice exposed to aerosols of respirable size. **(A)** Schematic of the TSS ^37^, including (1) Blaustein atomizer module, (2) syringe pump, (3) mouse restrainer and (4) scattered-light aerosol detector. **(B)** Lung CFU 24 h after a 20-min exposure of BALB/c mice to aerosols of the indicated knockout (KO) strains relative to the CFU for wild type H37Rv (Wt-rv) tested on the same day. The CFU level for Wt-rv across all experiments averaged 101.0 ± 20.2 (mean ± SD). **(C)** Restoration of wild type phenotype upon complementation of *rv0805* KO (*rv0805 comp*) and *rv1339* (*rv1339 comp*). For **(B)** and **(C)**, data were analyzed by one-way ANOVA and Dunnett’s post-test to correct for multiple comparisons using Mtb H37Rv as a reference. \*\**p*<0.01; \*\*\**p*< 0.001; \*\*\*\**p*< 0.0001.

To test whether the CFU deficit observed at 24 hours was an effect of the transmission model’s sequential stages, we infected mice with wild type and *rv0805* knockout strains harvested directly from stage 0 (7H9 medium under 21% O_2_, 5% CO_2_) and observed no CFU deficit in the *rv0805* knockout strain **(fig. S12)**. Thus, the transmission deficit of the *rv080*5 knockout was dependent on the conditions through which the bacteria had passed before they were aerosolized.

### Stage-dependent variations in cAMP

We were intrigued to see that both of Mtb’s known cAMP phosphodiesterases (*rv0805* and *rv1339*) were among the genes required for Mtb to survive stage 4 of modeled transmission. In retrospect, genes encoding 11 of Mtb’s 15 known adenylyl cyclases ^38^ were candidates for supporting Mtb’s survival through stages 1, 2 and/or 3, and one adenylyl cyclase (encoded by *rv3645*) appeared to be required for Mtb to survive all 4 stages (**fig. S13**). Prompted by this observation, we monitored Mtb’s cAMP levels through the first 4 stages using both a fluorescent reporter (**Fig. 6A**) and ELISA (**Fig. 6B).** For a fluorescent reporter, we adapted the GFP-based sensor of Wang et al ^39^ for optimal expression in Mtb and confirmed its responsiveness to changes in cAMP levels (**fig. S14**), then further modified the reporter for constitutive expression of mCherry (**fig. S15**). The fluorescent reporter and ELISA both revealed a marked decrease in cAMP content of wild type Mtb in stages 2 and 4. At stage 4, cAMP levels reach significantly higher concentrations in the *rv0805* knockout than in the *rv1339* knockout. By comparison, during stages 1 and 2, the *rv1339* knockout exhibited higher cAMP levels than the *rv0805* knockout **(Fig 6B & fig S16A, B).** No significant changes were observed in stage 3 for either strain. These results suggest that Rv0805 is the primary regulator of cAMP at stage 4, whereas Rv1339 mainly functions during stages 1 and 2, while also influencing levels at stage 4.

**Figure 6.**
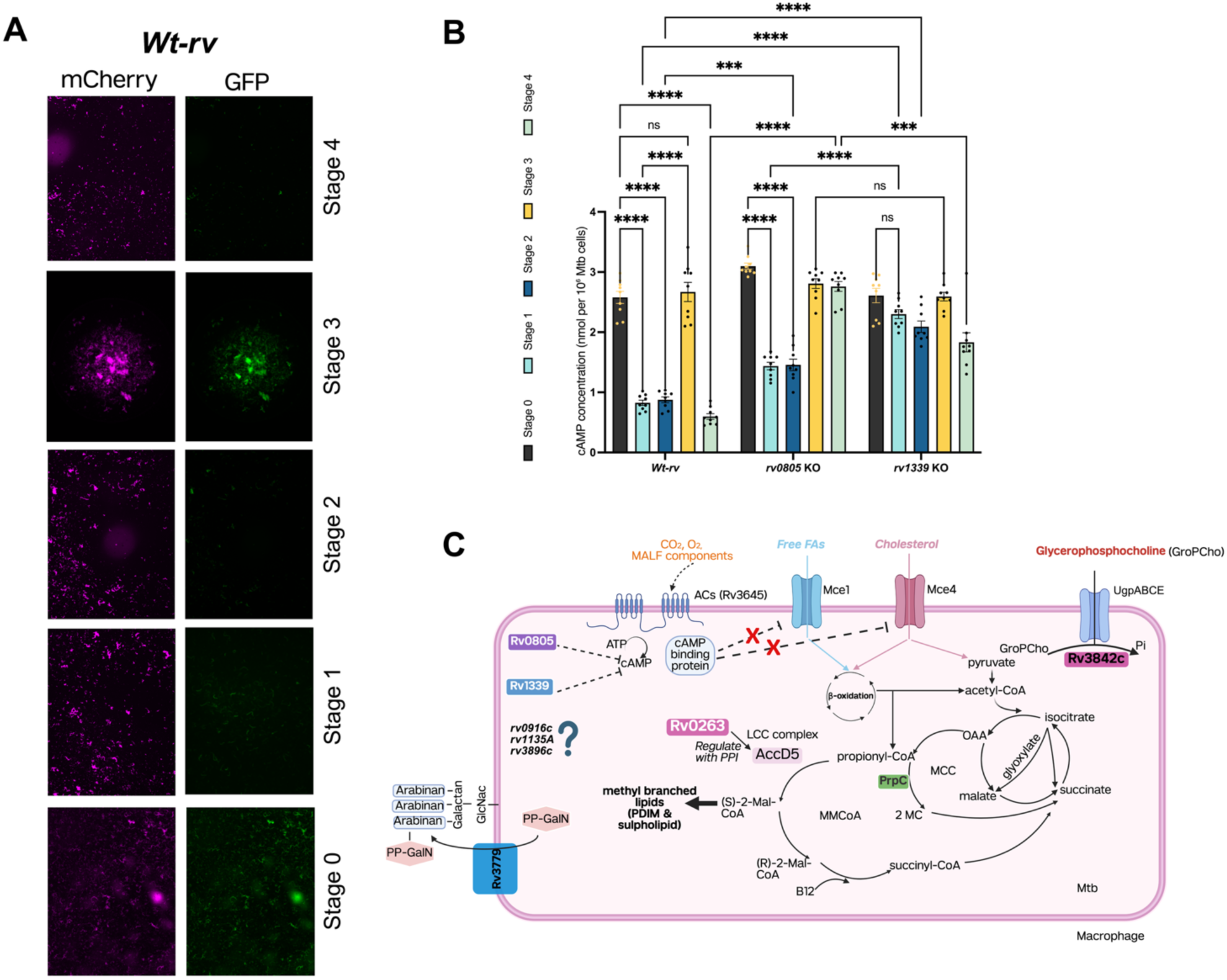
Cyclic AMP dynamics across modeled stages of transmission in wild-type and knock out strains of Mtb. **(A)** Fluorescence microscopy of Mtb cultures during stages 0 – 4. The bacteria constitutively express mCherry (magenta, top) as a control for the total number of bacteria, while GFP expression (green, bottom) reports high intracellular cAMP levels. Notable depletion of the GFP signal is observed in stages 1 and 2, with a transient recovery in stage 3 **(B)** Total cAMP concentrations measured across stages of the model for for Wt-rv and KO strains. Results are means ± SEM from 3 independent experiments. Significance is indicated by asterisks (*p < 0.05, **p < 0.01, ***p < 0.001; ns, non-significant). Statistical significance was determined by two-way ANOVA followed by Tukey’s multiple comparisons test. **(C)** Proposed interactions among some of the genes that Mtb requires for survival in the pulmonary alveolar environment after modeled aerosol transmission. PPI, protein-protein interactions

To understand what factors drove these changes in Mtb’s cAMP level, we first held the medium constant (MAF) and varied the levels of O₂ and CO₂ as applied in stage 1 (0.2% O₂, 5% CO₂), stage 2 (10% O₂, 5% CO₂), stage 3 (21% O₂, 0.04% CO₂) and stages 4 and 5 (21% O₂, 5% CO₂). We also tested a nonphysiological gas combination (0.2% O₂, 0.04% CO₂) **(fig. S17A**). The results suggested that Mtb incubated in MAF senses the level of both O_2_ and CO_2_ in regulating its cAMP levels, with the atmosphere of stage 1 (hypoxic lung lesion) producing the lowest level and the atmosphere of stage 3 (air) producing the highest level (**fig. S17B**).

Comparing cAMP levels of log phase Mtb in 7H9 medium in 21% O₂, 5% CO₂ to Mtb in MAF in the same atmosphere suggested that MAF itself also leads Mtb to decrease its levels of cAMP (**fig. S17B**).

Given this evidence that the liquid phase can also affect Mtb’s cAMP levels, we next tested the effect of MALF components on cAMP levels while holding the atmosphere constant at 21% O₂, 5% CO₂ as we rehydrated desiccated Mtb from stage 3 **(fig. S18A)**. Compared to rehydration in MEM-α or 7H9, which both maintained high cAMP concentrations (∼2.9 nmol/10^8^ Mtb), it emerged that surfactant proteins were chiefly responsible for the decrease in cAMP to ∼1 nmol/10^8^ Mtb seen when Mtb was rehydrated in MALF (**fig. S18B**).

In sum, Mtb regulates its levels of cAMP in response to both its gaseous and liquid environments throughout the modeled stages of transmission.

## Discussion

A key reason why about half of Mtb’s genes have unknown functions and over three-quarters of them are considered nonessential may be the experimental difficulty of seeking phenotypes in conditions that recapitulate the daunting sequence of environmental conditions that Mtb endures to complete its life cycle. There is likely considerable evolutionary pressure on Mtb’s genome from the need to pass from hypoxic lesions in a host whose potentially imminent mortality threatens Mtb’s ability to persist as a species, through the air and into the pulmonary alveoli of a new host, where Mtb encounters the lung’s innate antibacterial defenses ^40^.

To help close this knowledge gap, we put a genome wide library of Mtb knock-down strains through a model of 4 sequential stages of Mtb transmission. This required developing two different liquid phases, based on all the information we were able to gather about two distinct anatomic sites that are each lipid-rich but rich in different lipids: necrotic human lung and healthy human ALF. In our earlier report ^9^, we used MAF to mimic material spilling from a necrotic pulmonary lesion into the bronchial tree before it is expelled into the air in small droplets ^6,41^. For the present report, we made a comparable effort to model ALF so we could identify genes Mtb that appears to require for recovery from its airborne journey.

To use BALF itself for the bulk of these studies would have required BALF from over 90 donors. Given the substantial inter-donor variability in the impact of BALF on Mtb ^42^, we would have had to pool BALF from all donors before launching the screen. The cost, availability of volunteers, and time available for research protocols in the bronchoscopy suite made this prohibitive.

Instead, we formulated MALF based on multiple literature reports on the composition of BALF ^11–15,17–29^. Complicating this effort, each publication reported the concentration of different components of ALF. We combined the reported components at the reported concentrations. As a result, MALF is substantially more complex than other reported versions of simulated ALF. Perhaps because of differences in donor age ^43^, sex ^44^, residence in areas with or without substantial air pollution ^45^, and other variables ^46^, we confirmed reports ^31,33^ of the marked variability in Mtb’s transcriptomic response to BALF from different donors. This complicated the use of BALF to benchmark MALF. Nonetheless, among the genes consistently increased in expression on transfer of Mtb from 7H9 to rcBALF, 93% were also increased in Mtb upon transfer to MALF. When the Mtb had been passed through the first 3 stages of modeled transmission and then rehydrated, the transcriptomes of Mtb in MALF and in rcBALF were indistinguishable. With respect to Mtb coming from conventional culture into rcBALF, we observed many more changes in gene expression than others have described ^42^. This may be because we modified a standard method to compensate for the dilution of ALF during bronchoalveolar lavage.

Mtb coming out of conventional log phase culture in 7H9 under 21% O_2_, 5% CO_2_ (stage 0) showed no survival defect when exposed to MALF or rcBALF for 18 hours, the longest time we tested, consistent with the findings of Allué-Guardia et al for Mtb in BALF ^42^. However, when Mtb was passed through the initial three stages of modeled transmission and then exposed to MALF or rcBALF, survival defects emerged sometime between 3 hours and 6 hours. This observation suggests that aerosol-experienced Mtb must escape from ALF within 6 hours to maintain full viability. This is far longer than the 10 minutes it took alveolar macrophages to ingest adenoviruses delivered to the lung ^34^. Early after infection, alveolar macrophages provide a permissive replicative niche for Mtb ^47^, but our results suggest that even survival in immunologically naïve alveolar macrophages may require some of the same genes that were required for Mtb’s successful resuscitation in MALF.

From a methodologic standpoint, our results demonstrate that a NOSD as low as 1 in the pooled, genome-wide CRISPRi library sometimes correctly flags genes that have a phenotype when the individual knockouts or knockdowns are constructed and tested. As a technological platform, the TSS offers a substantive advance in verisimilitude for studies of aerosol transmission of Mtb. However, the flight path from aerosol generator to mouse nose is only 42.5 cm and the aerosol particles are exposed to air for only ∼4.0 sec before inhalation. This contrasts with the hours that human-generated aerosols of comparable size can remain suspended and minimizes the extent and duration of stresses associated with desiccation. Thus, the TSS only recapitulates the briefest in a range of transmission events. Moreover, we do not know to what extent mouse ALF and mouse alveolar macrophages resemble their human counterparts.

From a biological perspective, our results provide striking examples of the condition-dependence of essentiality. More important, our findings enlarge the concept of conditional essentiality by demonstrating that a phenotype may not emerge on transition from standard laboratory conditions to a given set of pathophysiologically more relevant conditions unless the organism has first passed through other pathophysiologically relevant conditions. We term this dependence on sequential transitions “dynamic essentiality.” This may be distinct from the temporally dependent essentiality described by Theriault et al. ^48^. There, the essentiality of different sets of Mtb genes was evident in single cell suspensions of mouse lungs taken at different times after infection ^48^, without implication that conditional essentiality at a later time point depended on Mtb having first experienced the conditions at an earlier time point. The study of dynamic essentiality offers increased understanding of gene function across a microbial pathogen’s life cycle. In terms of translational implications, some of the genes identified here might encode a protein that could serve as a target for a drug that could protect heavily exposed people from contracting TB. Candidates for such prophylaxis in high-incidence areas might be certain health care workers, visitors, and family members of newly diagnosed TB patients. In high-incidence areas, in-patient capacity is often limited during the onset of therapy, leading to early discharge. Moreover, it is often not known whether the infecting strain is sensitive to the drugs prescribed for the patient. In such cases, patients on treatment who are quickly discharged into the community may be contagious for extended periods.

Besides the 35 genes on which we focused, we identified 713 additional genes as candidates for supporting Mtb’s survival in stage 4. Each of the 713 was also implicated in an earlier stage of transmission. Some of these genes might encode a target for a drug that could contribute to cure of an infected patient, which is of course another way to reduce transmission. To use such drugs prophylactically would risk selecting for resistance, impairing their value for therapy. Hence our interest in the potential for prophylaxis is greatest in the 21 genes required in stage 4 but not in earlier stages.

Exploration of this concept, along with greater insight into Mtb’s transmission biology, will require further research to probe the functions of genes nominated as contributing to Mtb’s survival upon landing in a new host. For 19 of the 35 genes featured here, the literature offers little insight.

However, 7 of the genes collectively offer a potential insight into Mtb’s intra-alveolar biology, as suggested in **Fig. 6C**: survival during stages 4 and 5 appears to depend in large part on genes regulating Mtb’s uptake, catabolism and synthesis of lipids, as influenced by cAMP. While Mtb’s survival during stages 1–3 of modeled transmission relied heavily on proteostatic mechanisms, including hydrophilin orthologs that may protect proteins from desiccation ^9^, the gene encoding the adenylyl cyclase Rv3645 was also implicated in Mtb’s survival in stages 1-3 ^9^ as well as stages 4 and 5. Of Mtb’s 15 known adenylyl cyclases, Rv3645 appears to be the only one that is non-redundant and conditionally essential for Mtb’s growth in the presence of long-chain fatty acids, leading to its designation as MacE for “major adenylyl cyclase enzyme” ^49^. Elevation of cAMP reduces Mtb’s uptake of long-chain fatty acids ^49^ and cholesterol ^50^, thereby decreasing toxicity from their metabolic byproducts, while also potentially limiting Mtb’s growth in host-like environments ^50^. As others have noted ^49,51^, this bidirectional impact of cAMP levels on Mtb in lipid-rich environments requires Mtb to closely regulate its level of cAMP. Mtb’s need to control cAMP levels directs attention to the two phosphodiesterases required for Mtb’s recovery in MALF. Rv1339, which is under positive selection in clinical isolates of Mtb ^52^, is Mtb’s chief degrader of 3’, 5’ cAMP ^53^. In contrast, Rv0805 is 50-fold ^53^ to 150-fold ^54^ more active on 2’, 3’ cAMP and 2’, 3’ cGMP than on 3’, 5’ cAMP, suggesting a role in degradation of RNA. Rv0805 is also required for optimal growth on cholesterol ^55^, apparently via its role in detoxification of the propionyl-CoA generated from cholesterol catabolism ^55,56^. In stage 3 (desiccation), Mtb’s survival was promoted by depletion of KstR1 (Rv3574) (**Table S5**), a repressor of cholesterol catabolism ^57,58^. This suggests there may be a stage-dependent pause in Mtb’s investment in cholesterol utilization during transition to a new host, followed by a resumption of cholesterol consumption.

Rv3842 is a phosphodiesterase whose substrates are the polar heads of Mtb’s glycerophospholipids ^59^. When inorganic phosphate (Pi) is limiting, Rv3842 can generate Pi from phospholipids to sustain Mtb’s growth ^60^. Rv3842 may also help Mtb generate Pi from the dipalmitoylphosphatidylcholine in surfactant as it is taken up along with Mtb into the phagosomes of alveolar macrophage ^60^.

Rv3484 (CpsA) may play several roles in protecting Mtb as the bacteria revive in MALF after aerobic desiccation in MAF and undergo phagocytosis in alveolar macrophages. CpsA blocks the assembly of the superoxide-producing NADPH oxidase (NOX2) on the mycobacterial phagosome ^61^. In the mouse, CpsA also enables Mtb to escape from alveolar macrophages as they gradually become pro-inflammatory; many intrapulmonary Mtb then reside instead in a subset of immigrant monocytes that appear to provide a more favorable environment ^47^. Moreover, CpsA is required for Mtb’s uptake of palmitate ^62^, calling attention once more to the role of lipid metabolism in Mtb’s intra-alveolar adaptation. In addition to these immune-evasive and metabolic functions, Mtb appears to prioritize structural integrity through specific glycosylation pathways. For example Rv3779 is a glycosyltransferase that transfers polyprenyl-linked galactosamine or N-acetylgalactosamine to the arabinogalactan of Mtb’s cell wall ^63^. This suggests that cell wall remodeling may be a critical factor for Mtb’s survival during exposure to alveolar surfactants. The proposed requirement for cell wall modification is complemented by likely changes in core metabolism, where Rv0263 may play a regulatory role. Though the function of Rv0263 is unknown, its homolog in *Mycobacterium smegmatis* forms a complex with the homolog of Rv0264. Those two proteins co-purify with the long chain acyl CoA carboxylase complex, where they decrease its activity as a propionyl CoA carboxylase ^64^.

In sum, Mtb genes potentially supporting early stages of infection in a new host can be functionally categorized into two groups. The genes *rv0805*, *rv1339*, *rv3842c*, *rv3779*, *rv0916*, *rv3484* and *rv3842* appear dedicated to cAMP homeostasis, lipid sensing, and phosphate acquisition, while *rv1131*, *rv2934*, *rv0743, rv2515* and *rv0263* cope with detoxification of propionyl-CoA and the other metabolic shifts required for intracellular survival in an environment different from what Mtb experienced in its previous host and en route to its new one.

To the extent that the model presented here recapitulates aerosol transmission in the wild, our results suggest that survival of Mtb as a species is likely to depend on a set of conditionally and dynamically essential, stage-dependent, mutually non-redundant genes. Some of them help aerosolized Mtb recover from in-flight desiccation and resist the antibacterial effects of ALF long enough to take refuge in alveolar macrophages and survive there.

## ACKNOWLEDGMENTS

We thank J. Xiang and Adrian Y Tan of the Weill Cornell Genomics Resources Core Facility for extensive sequencing, and analysis and Jordi B. Torrelles (Texas Biomedical Research Institute) for generous advice. We are deeply grateful to Jeremy Rock (Rockefeller University) for providing the CRISPRi library. We gratefully acknowledge Amrita Singh, Weill Cornell Medicine, for her input on the cAMP experimental plan. We thank Dirk Schnappinger and Arka Banerjee (Weill Cornell Medicine) for their guidance in constructing ORBIT knockout strains and CRISPRi, as well as for generously sharing plasmids. We gratefully acknowledge Eric Rubin and Mark Sullivan (Harvard T.H. Chan School of Public Health) for sharing protocols for establishing ALI cultures, and Michael Shiloh (UT Southwestern) for insightful discussions. We recognize the valuable contributions of Kohta Saito and Christopher David Brown (Weill Cornell Medicine) to various aspects of this work. We extend our appreciation to J. Roberts, C. Suh, A. Lee, E. Kaplan, A. Singh, Z. Yang, and D. Li (Weill Cornell Medicine) for their assistance in an experiment that required multiple personnel. We are indebted to Xi Kathy Zhou for her expertise in statistical analyses and critical discussions. We further thank Kristin Burns-Huang, Ben Gold, and Ruslana Bryk (Weill Cornell Medicine) for administrative support and helpful discussions.

## Funding

This research was supported by NIH grant P01AI159402 (KR, PI; CN, co-PI), the Abby and Howard P. Milstein Program in Chemical Biology and Translational Medicine (CN), Bill and Melinda Gates Foundation investment INV-070075 (BV) and grants to PRS and SM from the Potts Memorial Foundation and the TB Research Advancement Center (TRAC) grant P30 AI168433 from NIH (Daniel Fitzgerald, PI). The Department of Microbiology & Immunology at Weill Cornell Medicine is supported by the William Randolph Hearst Foundation.

## Author contributions

Conceptualization: PRS, CN. Experimentation: PRS, MG, SM, XJ, FT, MC. Provision of special materials: KYR, RJK, PLL, RGC, DK, BV. Data analysis: PRS, MG, AJ, MDeJ. Writing: PRS, CN. Reviewing and editing: CN

## Competing interests

The authors declare no competing interests

## Data and materials availability

The RNA and DNA sequencing data have been submitted to the NCBI (PRJNA1460275, PRJNA1209962). Python codes are deposited at https://github.com/richiam16/MAF_Nathan

## Notes

https://github.com/richiam16/MAF_Nathan

### Competing Interest Statement

The authors have declared no competing interest.

### Summary of Updates

The revision adds new coauthors and new data.

