## Supplemental information for "*Mycobacterium tuberculosis* Genes Needed for Intra-alveolar Resuscitation After Airborne Transmission"

[1]Department of Microbiology and Immunology, Weill Cornell Medicine, New York, NY 10065; [2]Hackensack Meridian Health Center for Discovery and Innovation, Nutley, NJ 07110; [3]Skaggs School of Pharmacy & Pharmaceutical Sciences, University of California San Diego, San Diego CA 92093-0021; [4]Department of Chemistry and Biochemistry, University of California San Diego, San Diego CA 92093-0021; [5]Laboratory of Host-Pathogen Biology, Rockefeller University, New York, NY 10021; [6]Department of Genetic Medicine, Weill Cornell Medicine, New York, NY 10065; [7]Department of Medicine, Weill Cornell Medicine, New York, NY 10065; [8]Department of Microbiology and Immunology, College of Veterinary Medicine, Cornell University, Ithaca, New York

[†]Contributed equally to this work

[‡]Present address: National Institute of Pharmaceutical Education and Research, Sector 67, S.A.S. Nagar-160062, Punjab, India

### Materials and methods

#### Bacterial strains and growth conditions

We grew *Mycobacterium tuberculosis* H37Rv (Mtb) in 7H9 broth (BD DIFCO™ Middlebrook, 271310) supplemented with 10% (v/v) OADC (oleic acid, bovine albumin, dextrose, catalase; BD Biosciences), 0.2% glycerol, and 0.02% tyloxapol at 37 °C with 5% $CO_2$ under ambient humidity as described [1]. For Mtb knockout strains, the medium included hygromycin (50 µg/mL) and kanamycin (20 µg/mL). The CRISPRi library described in Mishra et al. [1] was a kind gift of Jeremy Rock, Rockefeller University. Individual CRISPRi strains were cultured in the presence of 100 ng/mL anhydrotetracycline (ATc) and 20 µg/mL kanamycin. CFU were detected on 7H11 agar (Middlebrook Agar Base, M511) supplemented as described above, excluding tyloxapol.

#### Preparation of MALF

MALF was prepared using components identified in human BALF, with each constituent added at its respective identified concentration. The formulation included phosphatidylglycerol 2.32 µg/mL , phosphatidylethanolamine 269.28 ng/mL, phosphatidylinositol 709.24 ng/mL, phosphatidylcholine 7.78 µg/mL, sphingomyelin 243.96 ng/mL [2]; cholesterol 0.1 mg/mL, 1,2-dipalmitoyl-sn-glycerol-3-phospho-(rac)-1-glycerol (DPPC) 4.8 mg/mL, 1,2-dipalmitoyl-sn-glycero-3-phospho-(1'-rac-glycerol) (DPPG) 0.5 mg/mL (12); surfactant proteins (SP) SP-A 4.4 µg/mL [3], SP-B 0.74 µg/mL [4], SP-C 0.57 µg/mL [5], SP-D 0.42 µg/mL [3] , IgG 2.6 mg/mL [6], IgA 0.8 mg/mL [7], IgM 0.1 mg/mL [7], lysozyme [6], albumin 8.8 mg/mL [6], transferrin 1.5 mg/mL [6], alpha 1-antitrypsin 2 mg/mL [7]. Minimum Essential Medium (MEM)-α with nucleosides and without phenol red (Gibco, catalog no: 41061037) was used as the base medium to mimic the water-soluble metabolites in alveolar lining fluid (ALF). For the lipid mixture, phosphatidylglycerol, phosphatidylethanolamine, phosphatidylinositol, phosphatidylcholine, cholesterol, sphingomyelin, DPPC and DPPG were dissolved in chloroform and combined in amounts to give the above final concentrations in a small, round-bottom glass flask. Chloroform was removed using a rotary evaporator in a water bath at 55 °C for ~15–20 minutes. The dried lipid film was resuspended in prewarmed (55 °C) Hanks' Balanced Salt Solution (HBSS; Gibco, 14175095), followed by vigorous mixing for 20–30 minutes. To this lipid mixture, albumin, transferrin, immunoglobulins, α1-antitrypsin, lysozyme, surfactants, uric acid, glutathione, ascorbic acid, and the following hydrolases were added at concentrations specified in **Table S1**—acid phosphatase, α-mannosidase, α-galactosidase, β-galactosidase, α-glucosidase, β-glucosidase, α-xylosidase, α-fucosidase, arylsulfatase, fatty acid esterase-I, non-specific esterase, alkaline phosphatase, alkaline phosphodiesterase, phospholipase C, peroxidase, α-rhamnosidase, and fatty acid esterase-II [8]. None of the references used in guiding preparation of MALF mentioned urea. As a result, we inadvertently omitted urea in early experiments but for most experiments included urea at 5 mM. Direct comparison showed that results were similar with and without urea (**Figure S19**).

**Collection of BALF**

We obtained ALF from normal nonsmokers by fiberoptic bronchosopy [9]. After informed consent was obtained under an approved Weill Cornell Medicine Institutional Review

Board protocol, fentanyl and midazolam were administered for sedation and lidocaine was used as a topical anesthetic on the vocal cords and airways. A fiberoptic bronchoscope was advanced through a mouthpiece and through the vocal cords to a distal airway and gently wedged into a subsegmental bronchus. Sterile 0.9% saline solution (23 °C) was infused in 20 mL aliquots, using a syringe. After each aliquot, the fluid was immediately recovered into a trap using suction. The total volume used per site was 150 mL. Two sites were evaluated, with a total volume not exceeding 300 mL. The right middle lobe and lingula were the usual sites for lavage. Of the saline used for lavage, 45– 65% of the infused volume was recovered.

**Reconstitution of BALF**

We used a blood urea nitrogen (BUN) Colorimetric Detection Kit (Invitrogen™, EIABUN) to measure the concentration of urea in BALF and a blood sample collected from the BALF donor near the time of bronchoaveolar lavage. We divided the urea concentration in blood by the urea concentration in BALF to determine the extent to which saline had diluted ALF during lavage [10]. We centrifuged BALF at 300 x g for 5 min at 4 °C to pellet cells. The supernatant was concentrated using a 3-kDa molecular weight cutoff centrifugal filter (Amicon® Ultra Centrifugal Filter, Millipore, UFC9003) by centrifugation at 2682 x g at 4 °C until the volume was reduced by the factor necessary to reverse the fold dilution determined by the urea measurements. The concentrated BALF was transferred to dialysis tubing (Pur-A-Lyzer™ Mega Dialysis Kit, Sigma-Aldrich, PURG35010) and dialyzed against 2 liters of MEM-α with nucleosides and without phenol red (Gibco Thermo Fisher Scientific, 41061037) at 4 °C for 24 h, during which the dialysis medium was replaced three times. The dialysand was designated reconstituted BALF (rcBALF).

**Osmolality and pH measurements**

The osmolality of MALF and reconstituted BALF (rcBALF) was measured with a Micro-Osmometer model 3300 (Advanced Instruments). The pH of MALF, rcBALF, and 7H9 was measured using MQuant® 1.09535 pH strips.

**Nontoxicity of MALF for primary human lung cells**

Cells pelleted from BALF were seeded onto collagen-coated Transwell inserts and cultured for 4 days in RPMI (Gibco,11875093) supplemented with 10% FBS (HyClone™ SH30088.03). After 4 days, the culture medium was removed from both the apical and basal sides. To the basal chamber we added 0.8 mL of MALF. After 24 hrs cells were collected from the apical side in 200 mL of PBS and transferred to a 96-well plate for viability assay using a CellTiter-Glo® Luminescent Cell Viability Assay kit according to the manufacturer's instructions.

**In vitro model of sequential stresses of transmission**

Throughout, when we refer to the concentrations of $O_2$ and $CO_2$, the balance of the gas phase is $N_2$. Mtb cultures grown in 7H9 medium in air with 5% $CO_2$ to an OD580 of 1 were washed and resuspended in MAF. To mimic the closed cavity state, these cultures were incubated under 0.2% $O_2$ and 5% $CO_2$ for two weeks. To simulate the transition to an open cavity (cavitation and contact with large airways), cultures were then incubated for an additional two weeks under 10% $O_2$ and 5% $CO_2$. To model aerosol formation, 2-µL droplets of the culture were deposited onto a polystyrene surface and allowed to dry above Drierite (Fisher Scientific NC9979539) for 24 hours (relative humidity, < 5%). For the inhalation step, dried Mtb was rehydrated in 100 µL MALF in air with 5% $CO_2$ and CFUs determined at 0, 3, 6, 10 and 24 hours. As a control, one arm of the experiment involved rehydrating the desiccated sample in 7H9 medium instead of MALF and determining CFU at the same intervals. For CRISPRi screening in the 4th stage, the CRISPRi library pre-depleted in response to Atc for 5, 10 and 20 generations and brought to an OD540 of 0.8-1.0 was then carried through the 3 stages of the transmission model as described (10) using 2 weeks each for stages 1 and 2. For the 3rd stage, 2-µL droplets were dispensed into single well plates (CELLTREAT plates 229501) using an Integra Viaflo 384 automated pipette and allowed to desiccate in a box above Drierite for 24 hrs at RT. For the 4th stage, cells were collected from the desiccated 2-uL droplets on single-well plates by adding 15 mL of MEM-α with 0.02% tyloxapol per plate, incubating for 20 minutes, and scraping. The cells were pooled in 50 mL Falcon tubes and centrifuged at 3098 x g, for 10 minutes at RT. The cell pellet was resuspended in MALF in 3 replicates of 3 mL and incubated at 37 °C with 5% $CO_2$ in air for 18 hours, centrifuged (3098 x g, 10 minutes, RT) and stored at -80 °C for gDNA extraction, sequencing, and MAGeCK-based CRISPRi data analysis as described [1].

**Transcriptomic analyses**

Wild type Mtb cultures were passed through the first 3 stages of modeled transmission as described above, except that the desiccated droplets from some of the plates in stage

3 were scraped into 15 mL of 2.5 M guanidinium thiocyanate per single-well plate. The desiccated cells in other stage 3 plates were scraped into 15 mL of MEM-α with 0.02% tyloxapol, centrifuged and resuspended in MALF (200 µL per plate) or in rcBALF (~66 µL per plate) and incubated for 18 h at 37 °C in 5% $CO_2$ and 21% $O_2$. Samples were centrifuged (3098 x g, 10 minutes, RT) and the pellets mixed with 1 mL of Trizol and 0.5 mL of silica beads for bead beating through 6 cycles of 1 minute each at 400 rpm, with 2 minutes on ice in between. Tubes were centrifuged at 13,523 x g, 10 minutes, at 4 °C. The supernatant was transferred to new tubes, mixed with 400 µL of chloroform, mixed with an equal amount of ethanol and processed for RNA extraction following the Zymo Research kit instructions for RNA isolation. RNA was quantified by NanoDrop and residual DNA digested with DNase using the TURBO DNA-free™ Kit (AM1907).

RNA was also isolated from wild type Mtb that did not pass through the modeled stages of transmission but was collected from 7H9 broth in log phase growth under air with 5% $CO_2$ at an OD580 of 1. One set of cells (~11.5 mL for one replicate) was mixed with an equal volume of 5 M guanidinium isothiocyanate and centrifuged (3098 x g, 10 minutes, RT). The pellet was resuspended in 1 mL of Trizol with 0.5 mL of silica beads for RNA isolation as above. Two other sets of log-phase Mtb cells (OD 1, ~11.5 mL per replicate) were pelleted and washed twice with PBS containing tyloxapol. One set was resuspended in 3 mL of MALF and the other in 1 mL of rcBALF. After 18 h, an equal volume of 5 M guanidinium isothiocyanate was added to each and the samples were centrifuged (3098 x g, 10 minutes, RT). The resulting pellets from Mtb incubated in MALF or rcBALF were resuspended in 1 mL of Trizol with 0.5 mL of silica beads for RNA isolation and DNase treatment as above.

RNA concentrations from all samples (stages 0 - 4) were measured using a NanoDrop. The Weill Cornell Medicine (WCM) sequencing core prepared RNA-seq libraries using the Illumina Stranded Total RNA Prep with Ribo-Zero Plus kit, following the manufacturer's instructions. Sequencing was done on a NovaSeq 6000 with paired-end 100 bp reads. The raw data in BCL format were converted to FASTQ files and sorted by sample using bcl2fastq version 2.20. After trimming adaptor sequences from the RNA reads using Cutadapt (version 1.18), the reads were aligned to the Mtb H37Rv genome using Bowtie 2 (version 2.2.8). The number of reads for each gene was counted with HTSeq-count (version 0.11.2). We used the DESeq2 package to analyze gene expression, group genes with similar patterns, and perform principal component analysis. To find genes that were expressed differently between groups, we compared them using parametric tests based on a negative binomial model, which accounts for differences in data variability. We corrected the p-values for multiple testing using the Benjamini-Hochberg method.

**Mtb infection of alveolar macrophages at an air-liquid interface on pulmonary epithelium**

A model of pulmonary macrophages at an air-liquid interface (ALI) above pulmonary epithelial cells was based on the protocol of Sullivan et al. [11]. Cells from the immortalized human epithelial cell line NuLi-1 (from ATCC) were expanded in a T25 flask for 4–5 days in serum-free Airway Epithelial Cell Basal Medium (BEGM; ATCC PCS-300-030) with additives from the Bronchial Epithelial Cell Growth Kit (ATCC PCS-300-040) as described [11]. To prepare Transwell (Falcon® 353104) inserts for ALI culture, 100 µL of human collagen (Sigma- H4417, 50 ug/mL in PBS) was added to each well in 24-well plates (Falcon® 353504). After 24 h at 4 °C, we aspirated the PBS and added approximately 90,000 NuLi-1 cells per well, followed by addition of 0.8 mL of BEGM to the basal side and 0.5 mL of BEGM to the apical side. After 2–3 days, we removed medium from both sides and replaced the basal medium with 0.8 mL of ALI differentiation medium [11]. Every 1-2 days thereafter, we removed liquid from the apical surface and replaced the basal medium.

Epithelial cell differentiation was assessed by expression of MUC5AC and F-actin. Cells were fixed directly on Transwell inserts by adding 4% paraformaldehyde (Thermo Scientific 043368) in PBS for 1 hour at room temperature. The medium was replaced with PBS and the cells incubated another h at 4 °C before permeabilization with 500 µL of PBS containing 1 mM $CaCl_2$ (MilliporeSigma C4901), 1 mM $MgCl_2$ (MilliporeSigma, M8266) and 0.2% Triton X-100 (MilliporeSigma T8787). After 15 min at RT, the membranes were placed in microcentrifuge tubes, washed with PBST (PBS with 1 mM $MgCl_2$, 1 mM $CaCl_2$ and 0.1% Tween 20 (MilliporeSigma, P1379) and placed on a rocking platform. Phalloidin-iFluor 488 (Abcam, ab176753) diluted 1:1000 in PBS with 1% bovine serum albumin (MilliporeSigma A9647) was used to stain F actin over 2 h. To stain for MUC5AC, membranes were rocked for 2 h at RT in PBST with 1% bovine serum albumin and 10% goat serum (Abcam ab7481), incubated overnight at 4 °C with anti-MUC5AC antibody (Abcam, ab3649), washed with PBST, and stained for 2 h at room temperature with 100 µL of anti-mouse IgG AlexaFluor 594 secondary antibody (Abcam, ab150116,) diluted 1:200 in PBST with 1% bovine serum albumin and 10% goat serum. After staining, all membranes were washed with PBST and nuclei were stained by adding 2 drops of

NucBlue in PBS for 1 min at RT, followed by another PBS wash. The membranes were mounted on glass slides (VWR, 16004430). ProLong™ Gold Antifade Mountant (Invtrogen, P36930) was added and the membranes covered with 22 mm × 22 mm No. 1.5 thickness glass coverslips. For confocal images, Phalloidin-iFluor 488 was excited at 470 nm, AlexaFluor 594 at 555 nm, and NucBlue™ at 460 nm for viewing in a Keyence microscope (BZ-X800) with a 100x oil-immersion objective.

After 14 days, by which time the epithelial cells were differentiated, epithelial cell monolayer permeability was assessed by adding 200 µL of sodium fluorescein (50 µg/mL; 46960, Millipore Sigma) to the apical compartment of a 24-well Transwell plate at 0, 3, 6, 9, and 14 days of ALI culture. Collagen-coated cell-free wells served as controls. After incubation for 30 minutes, 200 µL of PBS was added to both the apical and basal compartments to recover the total liquid. The collected samples were transferred to a 96-well plate and fluorescence was measured at an excitation wavelength of 482 nm and emission wavelength 527 nm. Monolayer permeability was determined by comparing fluorescein levels in the basal and apical compartments, normalized to values from collagen-coated control wells.

To the differentiated epithelial cultures, we added 50,000 differentiated mouse alveolar macrophage cells (AMJ2-C8 CRL-2455™, from ATCC) to the apical surface of the epithelial layer. One day later we added Mtb at a multiplicity of infection (MOI) of 1 relative to the macrophages. After 3 h, defined as 0 h post infection, we replaced the apical medium 3 times with 200 µL of PBS to remove extracellular Mtb. In some wells, we determined the initial uptake of Mtb by lysing the cells with 200 µL of 0.5% TritonX-100 in PBS in the apical compartment, followed by serial dilution in the same medium and plating on 7H11 agar to determine CFU. When the infection involved a gene knockout strain, the agar included 50 μg/mL hygromycin and 25 μg/mL kanamycin. For other wells, the medium was replaced on both the apical and basal sides at 24 and 48 h. Cells were lysed at 72 hours to determine CFU.

In one control experiment, a monolayer culture of differentiated mouse alveolar macrophage cells was seeded at a density of 50,000 cells per well in a 96-well plate. The cells were infected at an MOI of 1 with Mtb strains previously grown under stage 0 conditions (7H9 medium, 21% $O_2$, 5% $CO_2$). Following the protocol above, the macrophages were lysed at 0 h and 72 h post-infection, and the lysates were plated on 7H11 agar for CFU count.

**Mtb infection of human monocyte-derived macrophages**

Under a WCM IRB approved protocol, venous blood was collected into heparinized tubes from healthy adult volunteers. Monocytes were isolated isolated by CD14 magnetic bead selection (MACS; Miltenyi Biotec) and differentiated into monocyte-derived macrophages (MDM) as described [12]. Following the establishment of the Air-Liquid Interface (ALI) with NULI-1 epithelial cells (as described above), 50,000 MDMs—stimulated with 0.5 ng/mL each of recombinant human GM-CSF (R&D Systems) and TNF (Biolegend)—were infected at day 11 with Mtb at an MOI of 1. The Mtb strains used had been passaged through stages 1 to 4 of modeled transmission. After a 3-hour infection period, the cells were washed three times with 200 µL of PBS to remove extracellular bacteria. Cells were then lysed according to the previously described protocol and the lysates were plated on 7H11 agar at 0 h and 72 h post-infection.

**Construction of Mtb knockout strains**

We used the ORBIT (Oligonucleotide-Mediated Recombineering Followed by Bxb1 Integrase Targeting) method to generate gene knockouts [13]. Single-stranded 100-bp oligonucleotides were designed as in Murphy et al. [13]. Cultures of Mtb carrying the plasmid pKM461 were induced with 500 ng/mL ATc for 24 hours. Electrocompetent cells were prepared as described [1] and the cells were electroporated with a mixture of the oligonucleotides and pKM464 as described by Murphy et al. [13]. The following day, the transformed cells were plated on 7H11 agar containing 20 µg/mL kanamycin and 50 µg/mL hygromycin. After 4 weeks, single colonies were picked and expanded for genomic DNA isolation [1]. Screening for knockouts was performed using both PCR-based confirmation (**fig. S20)** and whole genome sequencing at Plasmidsaurus (Portland, Oregon, USA). For genomic PCR, we used two primer sets. In primer set 1, forward and reverse primers (**table S4**) flanking the upstream and downstream regions of the target gene were expected to yield a product of approximately 3100 bp, corresponding to the insertion of the hygromycin/oriE cassette in place of the gene. In primer Set 2, a forward primer that was the same as for first set and a reverse primer (**table S4**) within the hygromycin cassette were expected to yield a ~600 bp product, confirming replacement of part of the gene with the hygromycin cassette. For the generation of the complementation construct, the constitutive promoter of *hsp60* was utilized, following the methodology established by Beites et al. 2021[14]. The target gene was synthesized (GenScript) and integrated into the final expression vector, pMCK-hsp60-gene, via Gateway LR Clonase reaction using plasmids pEN12A-Phsp60, pEN41A-T02, and pDE43n-MCgZq19. Following transformation into the respective knockout strains, transformants were selected on 7H10 agar supplemented with hygromycin (50 ug/mL) and Zeocin (25 ug/mL). Successful complementation was confirmed by PCR analysis **(fig. S21)** of isolated genomic DNA (primers listed in **table S4**) and further validated through whole-genome sequencing.

**Construction of CRISPRi knockdown strains**

Single guide RNA (sgRNA) primers were designed using Pebble, an online platform maintained by the Jeremy Rock Lab at Rockefeller University (https://pebble.rockefeller.edu/), by selecting the non-template strand. The selected top and bottom oligonucleotides were annealed and ligated into EspI-digested pIRL58 (Addgene catalog no. 166886). These constructs were confirmed by Sanger sequencing. The pIRL19 plasmid (200 ng), containing the integrase, was co-transformed with the sgRNA-pIRL58 construct (200 ng) into Mtb following the protocol in Mishra et al. [1].

**Mouse infections**

Ten-week-old female BALB/c mice (Charles River Laboratories) were kept in individually ventilated cages in groups of 5 and offered high-fiber rodent chow and water ad libitum. Wild type Mtb H37Rv, knockout and complemented strains were incubated for 2 weeks each under stage 1 and stage 2 conditions in MAF prior to infecting groups of 8-10 animals using the TSS system as described [15]. Briefly, individual mice were simultaneously inoculated by nose-only exposure to aerosolized Mtb for 20 min. During this period, an inoculum of approximately 1 × $10^7$ CFU was aerosolized, resulting in the deposition of ~100 CFU/mouse for Mtb H37Rv, which served as a reference strain. At 24 hours post-infection, lungs of euthaized mice were collected, homogenized in 1 mL of PBS/0.05% Tween and plated in their entirety onto agar. Colonies were counted after 3-4 weeks of incubation at 37°C. All procedures involving animals were reviewed and approved by the Hackensack Meridian Health institutional animal care and use committee.

**Fluorescence microscopic evaluation of cAMP levels**

To evaluate cAMP levels during various stages of transmission, Mtb was transformed with the pVV16-hsp60 plasmid containing a dual-reporter cAMP sensor (mCherry-hsp60 and G-Flamp-GFP). Mtb cultures were grown in 7H9 medium to an OD600 of 1.0, pelleted, and resuspended in MAF before incubation in the in vitro transmission model. At stage 0 (log phase in 7H9 in 21% $O_2$ and 5% $CO_2$) and stages 2, 3, and 4, cultures were transferred to a 96-well plate for imaging. Fluorescence microscopy was performed using a Cy3/TRITC filter to detect the constitutive mCherry-hsp60 signal and a BZ-X GFP filter to monitor the GFP-cAMP reporter expression, allowing for the visual assessment of changes in cAMP levels.

**cAMP quantification**

Approximately 1 × $10^8$ Mtb grown under the stated conditions were pelleted and supernatants were collected for quantification of extracellular level of cAMP. The cell pellets were washed with 1× PBS containing Tween (PBS + Tween). The cells were resuspended in 500 μL of the lysis buffer provided with the cAMP assay kit (ab138880) and lysed by bead beating for 3 cycles on ice. After centrifugation at 13,000 rpm for 10 min, the supernatant was passed through a 0.22 μm filter. Samples were adjusted to contain equal quantities of protein and cAMP levels measured using a fluorescence-based cAMP detection kit (Abcam) in a plate reader according to the manufacturer's instructions. Total cAMP levels were determined by combining the values of secreted and cell-retained cAMP.

**Table S1. Composition of MALF**

| **1- Proteins** | | |
|---|---|---|
| **Surfactans (SP)** | **Concentration µg /mL** | **CatLog#** |
| SP-A | 4.4 | Creative biomart SFTPA1-2445H |
| SP-D | 0.42 | Creative biomart SFTPD-238H |
| SP-B | 0.74 | Creative biomart SFTPB-2395H |
| SP-C | 0.57 | Creative biomart SFTPC-3850H |
| **Hydrolases** | | |
| Acid phosphatase (Acp) | 0.57 | Human, SigmaAG60 |
| α-galactosidase (α-GAL) | 7.5 | E. coli, SigmaG7163-60UN |
| β-galactosidase (β-GAL) | 0.23 | E. coli.,Sigma10105031001 |
| α-glucosidase (α-GLU) | 2.4 | Bacillus stearothermophilus,SigmaG3651-250UN |
| β-glucosidase (β-GLU)) | 5.66 | Almonds,SigmaG4511-250UN |
| α-xylosidase (α-Xyl) | 77.77 | Megazyme, SigmaE-AXSEC |
| α-fucosidase (α-Fuc) | 100 | E. coli, SigmaF5884-2UN |
| Arylsulfatase (ARSI) | 0.8 | Helix pomatia,SigmaS9626-5KU |
| Fatty acid esterase-I | 1.33 | Porcine pancreas,SigmaL3126-100G |
| Non-specific esterase (NSE) | 34.66 | Porcine liver, SigmaE3019-3.5KU |
| Alkaline phosphatase (ALP) | 11.41 | Human Placenta, Sigma524604-100U |
| Alkaline phosphodiesterase (APDase) | 10 | Crotalus adamanteus venom, SigmaP3243-1VL |
| Phospholipase C (PLC) | 0.21 | Bacillus cereus, SigmaP5542-25UN |
| Peroxidase (POD) | 1.26 | Horseradish,Sigma77332-100MG |
| α-rhamnosidase (α-RHA) | 1.15 | Prokaryote,MegazymeE-RHAMS |
| Fatty acid esterase-II (FAE-II) | 0.0051 | Porcine pancreas, SigmaL0382-100KU |
| α-mannosidase (α- MAN) | 1.2 | Jack bean, AgilentGKX-5010 |
| Lysozyme | 60 | chicken egg white, SigmaL6876 |
| **Immunoglobulins** | | |
| IgG | 2.6 mg/ml | Sigma I4506 |

| | | |
|---|---|---|
| IgA | 800 | MP SKU0855906 |
| IgM | 100 | Sigma I8260 |
| **Others** | | |
| Albumin | 8000.8 | MP SKU0219134905 |
| Transferrin | 1000.5 | Sigma T0665 |
| Alpha 1-antitrypsin | 2000 | Sigma A6150 |
| **2- Lipids** | | |
| Phosphatidylglycerol | 2.32 | AvantiResearch 841138 |
| Phosphatidylethanolamine | 0.269 | Sigma 1535744 |
| Phosphatidylinositol | 0.709 | Sigma 79403 |
| Phosphatidylcholine | 7.78 | Sigma 1535733 |
| Sphingomyelin | 0.2243 | Sigma1535733 |
| Cholesterol (Natural lipid) | 100 | Sigma 8667 |
| DPPC | 4000.8 | Sigma P0763 |
| DPPG | 500 | AvantiResearch 840455 |
| **3- Metabolites** | | |
| MEM- $\alpha$ | 12% v/v | Gibco 41061037 |
| Glutathione | 49.66 | Sigma G4251 |
| Uric acid | 16.05 | Sigma U2625 |

**Table S2**

**Three healthy volunteers for BALF collection**

| Age | Sex |
|---|---|
| 43 years | Male |
| 45 years | Female |
| 69 years | Female |

**Table S3. Genes identified as candidates for contributing to Mtb's survival in transition from stages 1-3 to stage 4**

| No. | Gene id | NOSD | No. | Gene id | NOSD |
|---|---|---|---|---|---|
| 1 | rv3596c:clpC1 | 8 | 26 | rv3805c:aftB | 5 |
| 2 | rv0482:murB | 7 | 27 | rv1094:desA2 | 5 |
| 3 | rv0014c:pknB | 7 | 28 | rv0638:secE1 | 5 |
| 4 | rv0227c:rv0227c | 7 | 29 | rv1708:rv1708 | 5 |
| 5 | rv0732:secY | 7 | 30 | rv0639:nusG | 5 |
| 6 | rv0430:rv0430 | 7 | 31 | rv3646c:topA | 5 |
| 7 | rv0206c:mmpL3 | 7 | 32 | rv2682c:dxs1 | 5 |
| 8 | rv3808c:glfT2 | 7 | 33 | rv2165c:rv2165c | 5 |
| 9 | rv2093c:tatC | 7 | 34 | rv0015c:pknA | 5 |
| 10 | rv1697:rv1697 | 7 | 35 | rv0543c:rv0543c | 5 |
| 11 | rv0668:rpoC | 7 | 36 | rv1830:rv1830 | 5 |
| 12 | rv2592c:ruvB | 6 | 37 | rv0002:dnaN | 5 |
| 13 | rv2927c:rv2927c | 6 | 38 | rv2783c:gpsI | 5 |
| 14 | rv3285:accA3 | 6 | 39 | rv1436:gap | 5 |
| 15 | rv3794:embA | 6 | 40 | rv3801c:fadD32 | 5 |
| 16 | rv1111c:rv1111c | 6 | 41 | rv3265c:wbbL1 | 5 |
| 17 | rv2182c:rv2182c | 6 | 42 | rv1480:rv1480 | 5 |
| 18 | rv0667:rpoB | 6 | 43 | rv3048c:nrdF2 | 5 |
| 19 | rv2925c:rnc | 6 | 44 | rv3809c:glf | 5 |
| 20 | rv3645:rv3645 | 6 | 45 | rv0383c:rv0383c | 5 |
| 21 | rv2200c:ctaC | 6 | 46 | rv3240c:secA1 | 5 |
| 22 | rv3043c:ctaD | 6 | 47 | rv0001:dnaA | 5 |
| 23 | rv1828:rv1828 | 6 | 48 | rv2150c:ftsZ | 5 |
| 24 | rv6043:tRNA-Ser41 | 5 | 49 | rv3246c:mtrA | 5 |
| 25 | rv1297:rho | 5 | 50 | rv0058:dnaB | 5 |

| No. | Gene id | NOSD | No. | Gene id | NOSD |
|---|---|---|---|---|---|
| 51 | rv1302:rfe | 5 | 86 | rv3302c:glpD2 | 4 |
| 52 | rv1223:htrA | 5 | 87 | rv2243:fabD | 4 |
| 53 | rv6009:tRNA-Lys10 | 4 | 88 | rv2926c:rv2926c | 4 |
| 54 | rv0652:rplL | 4 | 89 | rv3783:rfbD | 4 |
| 55 | rv0716:rplE | 4 | 90 | rv2201:asnB | 4 |
| 56 | rv2890c:rpsB | 4 | 91 | rv3051c:nrdE | 4 |
| 57 | rv0701:rplC | 4 | 92 | rv2195:qcrA | 4 |
| 58 | rv2748c:ftsK | 4 | 93 | rv1311:atpC | 4 |
| 59 | rv0384c:clpB | 4 | 94 | rv3778c:rv3778c | 4 |
| 60 | rv0060:rv0060 | 4 | 95 | rv2156c:murX | 4 |
| 61 | rv3915:rv3915 | 4 | 96 | rv3587c:rv3587c | 4 |
| 62 | rv2445c:ndkA | 4 | 97 | rv1310:atpD | 4 |
| 63 | rv0351:grpE | 4 | 98 | rv3909:rv3909 | 4 |
| 64 | rv0282:eccA3 | 4 | 99 | rv1224:tatB | 4 |
| 65 | rv0006:gyrA | 4 | 100 | rv1611:trpC | 4 |
| 66 | rv1547:dnaE1 | 4 | 101 | rv0338c:rv0338c | 4 |
| 67 | rv0337c:aspC | 4 | 102 | rv0808:purF | 4 |
| 68 | rv3709c:ask | 4 | 103 | rv1594:nadA | 4 |
| 69 | rv1326c:glgB | 4 | 104 | rv3583c:rv3583c | 4 |
| 70 | rv3795:embB | 4 | 105 | rv2145c:wag31 | 4 |
| 71 | rv1484:inhA | 4 | 106 | rv2846c:efpA | 4 |
| 72 | rv2539c:aroK | 4 | 107 | rv1477:ripA | 4 |
| 73 | rv1462:rv1462 | 4 | 108 | rv3457c:rpoA | 4 |
| 74 | rv3807c:rv3807c | 4 | 109 | rv3245c:mtrB | 4 |
| 75 | rv0005:gyrB | 4 | 110 | rv2196:qcrB | 4 |
| 76 | rv0431:rv0431 | 4 | 111 | rv3859c:gltB | 4 |
| 77 | rv2703:sigA | 4 | 112 | rv2361c:rv2361c | 4 |
| 78 | rv3260c:whiB2 | 4 | 113 | rv2524c:fas | 4 |
| 79 | rv2673:aftC | 4 | 114 | rv0903c:prrA | 4 |
| 80 | rv3277:rv3277 | 4 | 115 | rv3779:rv3779 | 3 |
| 81 | rv3918c:parA | 4 | 116 | rv2476c:gdh | 3 |
| 82 | rv2572c:aspS | 4 | 117 | rv0541c:rv0541c | 3 |
| 83 | rv6019:23S | 4 | 118 | rv3459c:rpsK | 3 |
| 84 | rv1423:whiA | 4 | 119 | rv2975A:rv2975A | 3 |
| 85 | rv3782:glfT1 | 4 | 120 | rv1824:rv1824 | 3 |

| | | | | | |
|---|---|---|---|---|---|
| 121 | rv3053c:nrdH | 3 | 156 | rv0283:eccB3 | 3 |
| 122 | rv1130:prpD | 3 | 157 | rv0041:leuS | 3 |
| 123 | rv2535c:pepQ | 3 | 158 | rv3034c:rv3034c | 3 |
| 124 | rv2157c:murF | 3 | 159 | rv2996c:serA1 | 3 |
| 125 | rv0707:rpsC | 3 | 160 | rv2448c:valS | 3 |
| 126 | rv0721:rpsE | 3 | 161 | rv1343c:lprD | 3 |
| 127 | rv3193c:rv3193c | 3 | 162 | rv3247c:tmk | 3 |
| 128 | rv6024:tRNA-Leu22 | 3 | 163 | rv3858c:gltD | 3 |
| 129 | rv1630:rpsA | 3 | 164 | rv3722c:rv3722c | 3 |
| 130 | rv0410c:pknG | 3 | 165 | rv1782:eccB5 | 3 |
| 131 | rv0226c:rv0226c | 3 | 166 | rv1177:fdxC | 3 |
| 132 | rv0290:eccD3 | 3 | 167 | rv1649:pheS | 3 |
| 133 | rv2219:rv2219 | 3 | 168 | rv6008:tRNA-Thr9 | 3 |
| 134 | rv1694:tlyA | 3 | 169 | rv1825:rv1825 | 3 |
| 135 | rv0718:rpsH | 3 | 170 | rv3101c:ftsX | 3 |
| 136 | rv1348:irtA | 3 | 171 | rv1304:atpB | 3 |
| 137 | rv3105c:prfB | 3 | 172 | rv1463:rv1463 | 3 |
| 138 | rv1699:pyrG | 3 | 173 | rv3336c:trpS | 3 |
| 139 | rv0702:rplD | 3 | 174 | rv1683:rv1683 | 3 |
| 140 | rv2970A:rv2970A | 3 | 175 | rv2555c:alaS | 3 |
| 141 | rv3464:rmlB | 3 | 176 | rv2229c:rv2229c | 3 |
| 142 | rv2457c:clpX | 3 | 177 | rv1476:rv1476 | 3 |
| 143 | rv0635:hadA | 3 | 178 | rv0703:rplW | 3 |
| 144 | rv6013:tRNA-Arg14 | 3 | 179 | rv3793:embC | 3 |
| 145 | rv0883c:rv0883c | 3 | 180 | rv2921c:ftsY | 3 |
| 146 | rv2754c:thyX | 3 | 181 | rv6023:tRNA-Pro21 | 3 |
| 147 | rv3581c:ispF | 3 | 182 | rv1159:pimE | 3 |
| 148 | rv2540c:aroF | 3 | 183 | rv0685:tuf | 3 |
| 149 | rv0710:rpsQ | 3 | 184 | rv1617:pykA | 3 |
| 150 | rv1408:rpe | 3 | 185 | rv2109c:prcA | 3 |
| 151 | rv3792:aftA | 3 | 186 | rv2580c:hisS | 3 |
| 152 | rv0705:rpsS | 3 | 187 | rv2614c:thrS | 3 |
| 153 | rv1609:trpE | 3 | 188 | rv2244:acpM | 3 |
| 154 | rv3608c:folP1 | 3 | 189 | rv3044:fecB | 3 |
| 155 | rv2881c:cdsA | 3 | 190 | rv2152c:murC | 3 |

| | | | | | |
|---|---|---|---|---|---|
| 191 | rv1684:rv1684 | 3 | 226 | rv0225:rv0225 | 3 |
| 192 | rv1382:rv1382 | 3 | 227 | rv3355c:rv3355c | 3 |
| 193 | rv0636:hadB | 3 | 228 | rv2174:mptA | 3 |
| 194 | rv3917c:parB | 3 | 229 | rv0810c:rv0810c | 3 |
| 195 | rv2245:kasA | 3 | 230 | rv1389:gmk | 3 |
| 196 | rv3580c:cysS1 | 3 | 231 | rv0236A:rv0236A | 3 |
| 197 | rv2971:rv2971 | 3 | 232 | rv1636:TB15.3 | 3 |
| 198 | rv2786c:ribF | 3 | 233 | rv3799c:accD4 | 3 |
| 199 | rv1406:fmt | 3 | 234 | rv3834c:serS | 3 |
| 200 | rv3040c:rv3040c | 3 | 235 | rv2916c:ffh | 3 |
| 201 | rv3030:rv3030 | 3 | 236 | rv1306:atpF | 3 |
| 202 | rv3708c:asd | 3 | 237 | rv0364:rv0364 | 3 |
| 203 | rv2246:kasB | 3 | 238 | rv3908:mutT4 | 3 |
| 204 | rv2992c:gltS | 3 | 239 | rv2581c:rv2581c | 3 |
| 205 | rv1024:rv1024 | 3 | 240 | rv0526:rv0526 | 3 |
| 206 | rv3303c:lpdA | 3 | 241 | rv0357c:purA | 3 |
| 207 | rv1613:trpA | 3 | 242 | rv2845c:proS | 3 |
| 208 | rv0020c:fhaA | 3 | 243 | rv1481:rv1481 | 3 |
| 209 | rv0684:fusA1 | 3 | 244 | rv1110:lytB2 | 3 |
| 210 | rv3035:rv3035 | 3 | 245 | rv2610c:pimA | 3 |
| 211 | rv0236c:aftD | 3 | 246 | rv3604c:rv3604c | 3 |
| 212 | rv0777:purB | 3 | 247 | rv3790:dprE1 | 3 |
| 213 | rv0993:galU | 3 | 248 | rv3669:rv3669 | 3 |
| 214 | rv0286:PPE4 | 3 | 249 | rv1380:pyrB | 3 |
| 215 | rv2155c:murD | 3 | 250 | rv2256c:rv2256c | 3 |
| 216 | rv2247:accD6 | 3 | 251 | rv3410c:guaB3 | 3 |
| 217 | rv3264c:manB | 3 | 252 | rv1440:secG | 3 |
| 218 | rv1695:ppnK | 3 | 253 | rv3411c:guaB2 | 3 |
| 219 | rv0189c:ilvD | 3 | 254 | rv2121c:hisG | 3 |
| 220 | rv3764c:tcrY | 3 | 255 | rv1650:pheT | 3 |
| 221 | rv3436c:glmS | 3 | 256 | rv2968c:rv2968c | 3 |
| 222 | rv1483:fabG1 | 3 | 257 | rv0529:ccsA | 3 |
| 223 | rv1479:moxR1 | 3 | 258 | rv3781:rfbE | 3 |
| 224 | rv0824c:desA1 | 3 | 259 | rv0884c:serC | 3 |
| 225 | rv3800c:pks13 | 3 | 260 | rv0126:treS | 3 |

| | | | | | |
|---|---|---|---|---|---|
| 261 | rv3248c:sahH | 3 | 296 | rv6005:tRNA-Thr6 | 2 |
| 262 | rv2147c:rv2147c | 3 | 297 | rv0641:rplA | 2 |
| 263 | rv3257c:pmmA | 3 | 298 | rv3337:rv3337 | 2 |
| 264 | rv1166:lpqW | 3 | 299 | rv0936:pstA2 | 2 |
| 265 | rv2198c:mmpS3 | 3 | 300 | rv1458c:rv1458c | 2 |
| 266 | rv1451:ctaB | 3 | 301 | rv3814c:rv3814c | 2 |
| 267 | rv2612c:pgsA1 | 3 | 302 | rv1539:lspA | 2 |
| 268 | rv0440:groEL2 | 3 | 303 | rv2593c:ruvA | 2 |
| 269 | rv3244c:lpqB | 3 | 304 | rv3356c:folD | 2 |
| 270 | rv1642:rpmI | 3 | 305 | rv6041:tRNA-Thr39 | 2 |
| 271 | rv1309:atpG | 3 | 306 | rv6018:16S | 2 |
| 272 | rv3280:accD5 | 3 | 307 | rv2477c:rv2477c | 2 |
| 273 | rv3806c:ubiA | 3 | 308 | rv2984:ppk1 | 2 |
| 274 | rv2110c:prcB | 3 | 309 | rv0955:rv0955 | 2 |
| 275 | rv3258c:rv3258c | 3 | 310 | rv0017c:rodA | 2 |
| 276 | rv0956:purN | 3 | 311 | rv3712:rv3712 | 2 |
| 277 | rv2969c:rv2969c | 3 | 312 | rv2839c:infB | 2 |
| 278 | rv2194:qcrC | 3 | 313 | rv1565c:rv1565c | 2 |
| 279 | rv1308:atpA | 3 | 314 | rv0647c:rv0647c | 2 |
| 280 | rv2193:ctaE | 3 | 315 | rv3125c:PPE49 | 2 |
| 281 | rv3913:trxB2 | 3 | 316 | rv3672c:rv3672c | 2 |
| 282 | rv3370c:dnaE2 | 2 | 317 | rv2843:rv2843 | 2 |
| 283 | rv2517c:rv2517c | 2 | 318 | rv3423c:alr | 2 |
| 284 | rv0450c:mmpL4 | 2 | 319 | rv2008c:rv2008c | 2 |
| 285 | rv1056:rv1056 | 2 | 320 | rv2218:lipA | 2 |
| 286 | rv1770:rv1770 | 2 | 321 | rv0528:rv0528 | 2 |
| 287 | rv3682:ponA2 | 2 | 322 | rv6035:tRNA-Cys33 | 2 |
| 288 | rv1012:rv1012 | 2 | 323 | rv3910:rv3910 | 2 |
| 289 | rv1783:eccC5 | 2 | 324 | rv3012c:gatC | 2 |
| 290 | rv2238c:ahpE | 2 | 325 | rv3443c:rplM | 2 |
| 291 | rv2551c:rv2551c | 2 | 326 | rv2163c:pbpB | 2 |
| 292 | rv2909c:rpsP | 2 | 327 | rv1222:rseA | 2 |
| 293 | rv0047c:rv0047c | 2 | 328 | rv0347:rv0347 | 2 |
| 294 | rv1536:ileS | 2 | 329 | rv1221:sigE | 2 |
| 295 | rv1845c:blaR | 2 | 330 | rv6046:tRNA-Ser44 | 2 |

| | | | | | |
|---|---|---|---|---|---|
| 331 | rv0637:hadC | 2 | 366 | rv0207c:rv0207c | 2 |
| 332 | rv2343c:dnaG | 2 | 367 | rv3276c:purK | 2 |
| 333 | rv1604:impA | 2 | 368 | rv0003:recF | 2 |
| 334 | rv0709:rpmC | 2 | 369 | rv1017c:prsA | 2 |
| 335 | rv3916c:rv3916c | 2 | 370 | rv1292:argS | 2 |
| 336 | rv0334:rmlA | 2 | 371 | rv2169c:rv2169c | 2 |
| 337 | rv1294:thrA | 2 | 372 | rv3301c:phoY1 | 2 |
| 338 | rv1437:pgk | 2 | 373 | rv2869c:rip | 2 |
| 339 | rv0788:purQ | 2 | 374 | rv2215:dlaT | 2 |
| 340 | rv1456c:rv1456c | 2 | 375 | rv0046c:ino1 | 2 |
| 341 | rv2904c:rplS | 2 | 376 | rv0640:rplK | 2 |
| 342 | rv0683:rpsG | 2 | 377 | rv1449c:tkt | 2 |
| 343 | rv2870c:dxr | 2 | 378 | rv0706:rplV | 2 |
| 344 | rv2190c:rv2190c | 2 | 379 | rv0284:eccC3 | 2 |
| 345 | rv3236c:rv3236c | 2 | 380 | rv2516c:rv2516c | 2 |
| 346 | rv3456c:rplQ | 2 | 381 | rv1315:murA | 2 |
| 347 | rv1211:rv1211 | 2 | 382 | rv1018c:glmU | 2 |
| 348 | rv1641:infC | 2 | 383 | rv1378c:rv1378c | 2 |
| 349 | rv2158c:murE | 2 | 384 | rv1689:tyrS | 2 |
| 350 | rv1295:thrC | 2 | 385 | rv2112c:dop | 2 |
| 351 | rv0527:ccdA | 2 | 386 | rv1099c:glpX | 2 |
| 352 | rv0205:rv0205 | 2 | 387 | rv0524:hemL | 2 |
| 353 | rv3279c:birA | 2 | 388 | rv1392:metK | 2 |
| 354 | rv1410c:rv1410c | 2 | 389 | rv0719:rplF | 2 |
| 355 | rv0408:pta | 2 | 390 | rv1710:scpB | 2 |
| 356 | rv1610:rv1610 | 2 | 391 | rv1459c:rv1459c | 2 |
| 357 | rv3462c:infA | 2 | 392 | rv1713:engA | 2 |
| 358 | rv3804c:fbpA | 2 | 393 | rv1388:mihF | 2 |
| 359 | rv2678c:hemE | 2 | 394 | rv2178c:aroG | 2 |
| 360 | rv2228c:rv2228c | 2 | 395 | rv6000:tRNA-Ile1 | 2 |
| 361 | rv2837c:rv2837c | 2 | 396 | rv3609c:folE | 2 |
| 362 | rv3458c:rpsD | 2 | 397 | rv6047:tRNA-Ser45 | 2 |
| 363 | rv3221c:TB7.3 | 2 | 398 | rv1602:hisH | 2 |
| 364 | rv6042:tRNA-Pro40 | 2 | 399 | rv1300:hemK | 2 |
| 365 | rv3791:dprE2 | 2 | 400 | rv3924c:rpmH | 2 |

| | | | | | |
|---|---|---|---|---|---|
| 401 | rv0642c:mmaA4 | 2 | 436 | rv2204c:rv2204c | 2 |
| 402 | rv0455c:rv0455c | 2 | 437 | rv0896:gltA2 | 2 |
| 403 | rv2552c:aroE | 2 | 438 | rv2965c:kdtB | 2 |
| 404 | rv1293:lysA | 2 | 439 | rv1461:rv1461 | 2 |
| 405 | rv2676c:rv2676c | 2 | 440 | rv1023:eno | 2 |
| 406 | rv2357c:glyS | 2 | 441 | rv2461c:clpP1 | 2 |
| 407 | rv6007:tRNA-Trp8 | 2 | 442 | rv2981c:ddlA | 2 |
| 408 | rv6004:tRNA-Tyr5 | 2 | 443 | rv0204c:rv0204c | 2 |
| 409 | rv1390:rpoZ | 2 | 444 | rv1381:pyrC | 2 |
| 410 | rv6017:tRNA-Arg18 | 2 | 445 | rv3598c:lysS | 2 |
| 411 | rv2166c:rv2166c | 2 | 446 | rv1415:ribA2 | 2 |
| 412 | rv3721c:dnaZX | 2 | 447 | rv1307:atpH | 2 |
| 413 | rv3676:crp | 2 | 448 | rv2611c:rv2611c | 2 |
| 414 | rv2518c:ldtB | 2 | 449 | rv1303:rv1303 | 2 |
| 415 | rv1086:rv1086 | 2 | 450 | rv1338:murI | 2 |
| 416 | rv6040:tRNA-Met38 | 2 | 451 | rv3029c:fixA | 2 |
| 417 | rv2440c:obg | 2 | 452 | rv1208:gpgS | 2 |
| 418 | rv2111c:pup | 2 | 453 | rv1379:pyrR | 2 |
| 419 | rv1327c:glgE | 2 | 454 | rv0772:purD | 2 |
| 420 | rv3256c:rv3256c | 2 | 455 | rv0004:rv0004 | 2 |
| 421 | rv2553c:rv2553c | 2 | 456 | rv0016c:pbpA | 2 |
| 422 | rv2548A:rv2548A | 2 | 457 | rv3227:aroA | 2 |
| 423 | rv3275c:purE | 2 | 458 | rv6038:tRNA-Gln36 | 2 |
| 424 | rv0957:purH | 2 | 459 | rv2439c:proB | 2 |
| 425 | rv3418c:groES | 2 | 460 | rv0350:dnaK | 2 |
| 426 | rv2795c:rv2795c | 2 | 461 | rv2421c:nadD | 2 |
| 427 | rv6002:tRNA-Leu3 | 2 | 462 | rv2538c:aroB | 2 |
| 428 | rv6037:tRNA-Glu35 | 2 | 463 | rv2613c:rv2613c | 2 |
| 429 | rv2868c:gcpE | 2 | 464 | rv2207:cobT | 2 |
| 430 | rv3634c:galE1 | 2 | 465 | rv2122c:hisE | 2 |
| 431 | rv2401A:rv2401A | 2 | 466 | rv2220:glnA1 | 2 |
| 432 | rv2753c:dapA | 2 | 467 | rv1827:garA | 2 |
| 433 | rv1612:trpB | 2 | 468 | rv1007c:metS | 2 |
| 434 | rv2153c:murG | 2 | 469 | rv3396c:guaA | 2 |
| 435 | rv3610c:ftsH | 2 | 470 | rv3624c:hpt | 2 |

| | | | | | |
|---|---|---|---|---|---|
| 471 | rv2094c:tatA | 2 | 506 | Empty | 1 |
| 472 | rv3372:otsB2 | 2 | 507 | rv1427c:fadD12 | 1 |
| 473 | rv2460c:clpP2 | 2 | 508 | rv0743c:rv0743c | 1 |
| 474 | rv1339:rv1339 | 1 | 509 | rv2515c:rv2515c | 1 |
| 475 | rv0556:rv0556 | 1 | 510 | rv2953:rv2953 | 1 |
| 476 | rv2144c:rv2144c | 1 | 511 | rv0916c:PE7 | 1 |
| 477 | rv3156:nuoL | 1 | 512 | rv3896c:rv3896c | 1 |
| 478 | rv3196A:rv3196A | 1 | 513 | rv3882c:eccE1 | 1 |
| 479 | rv2287:yjcE | 1 | 514 | rv3567c:hsaB | 1 |
| 480 | rv1129c:rv1129c | 1 | 515 | rv0023:rv0023 | 1 |
| 481 | rv1131:prpC | 1 | 516 | rv3696c:glpK | 1 |
| 482 | rv3741c:rv3741c | 1 | 517 | rv3629c:rv3629c | 1 |
| 483 | rv1629:polA | 1 | 518 | rv3294c:rv3294c | 1 |
| 484 | rv3484:cpsA | 1 | 519 | rv1430:PE16 | 1 |
| 485 | rv2902c:rnhB | 1 | 520 | rv0007:rv0007 | 1 |
| 486 | rv3842c:glpQ1 | 1 | 521 | rv3616c:espA | 1 |
| 487 | rv3441c:mrsA | 1 | 522 | rv0099:fadD10 | 1 |
| 488 | rv0456c:echA2 | 1 | 523 | rv2377c:mbtH | 1 |
| 489 | rv0402c:mmpL1 | 1 | 524 | rv3140:fadE23 | 1 |
| 490 | rv1085c:rv1085c | 1 | 525 | rv0922:rv0922 | 1 |
| 491 | rv1653:argJ | 1 | 526 | rv1554:frdC | 1 |
| 492 | rv2171:lppM | 1 | 527 | rv3169:rv3169 | 1 |
| 493 | rv0263c:rv0263c | 1 | 528 | rv2081c:rv2081c | 1 |
| 494 | rv2002:fabG3 | 1 | 588 | rv6044:tRNA-Ser42 | 1 |
| 495 | rv2103c:vapC37 | 1 | 589 | rv0651:rplJ | 1 |
| 496 | rv2970c:lipN | 1 | 590 | rv6025:tRNA-Val23 | 1 |
| 497 | rv3268:rv3268 | 1 | 591 | rv1793:esxN | 1 |
| 498 | rv1135A:rv1135A | 1 | 592 | rv0899:ompA | 1 |
| 499 | rv3479:rv3479 | 1 | 593 | rv3032:rv3032 | 1 |
| 500 | rv3206c:moeB1 | 1 | 594 | rv1083:rv1083 | 1 |
| 501 | rv2482c:plsB2 | 1 | 595 | rv1201c:dapD | 1 |
| 502 | rv2934:ppsD | 1 | 596 | rv0986:rv0986 | 1 |
| 503 | rv1888c:rv1888c | 1 | 597 | rv0906:rv0906 | 1 |
| 504 | rv3606c:folK | 1 | 598 | rv1601:hisB | 1 |
| 505 | rv0805:rv0805 | 1 | 599 | rv2151c:ftsQ | 1 |

| | | | | | |
|---|---|---|---|---|---|
| 600 | rv2986c:hupB | 1 | 635 | rv3665c:dppB | 1 |
| 601 | rv3470c:ilvB2 | 1 | 636 | rv3670:ephE | 1 |
| 602 | rv0458:rv0458 | 1 | 637 | rv3346c:rv3346c | 1 |
| 603 | rv0032:bioF2 | 1 | 638 | rv3100c:smpB | 1 |
| 604 | rv3345c:PE-PGRS50 | 1 | 639 | rv2230c:rv2230c | 1 |
| 605 | rv3340:metC | 1 | 640 | rv0011c:rv0011c | 1 |
| 606 | rv1794:rv1794 | 1 | 641 | rv0412c:rv0412c | 1 |
| 607 | rv2958c:rv2958c | 1 | 642 | rv2441c:rpmA | 1 |
| 608 | rv3854c:ethA | 1 | 643 | rv3635:rv3635 | 1 |
| 609 | rv2609c:rv2609c | 1 | 644 | rv2149c:yfiH | 1 |
| 610 | rv0229Ac:rv0229Ac | 1 | 645 | rv0294:tam | 1 |
| 611 | rv2483c:plsC | 1 | 646 | rv1661:pks7 | 1 |
| 612 | rv3001c:ilvC | 1 | 647 | rv2903c:lepB | 1 |
| 613 | rv3267:rv3267 | 1 | 648 | rv3627c:rv3627c | 1 |
| 614 | rv2362c:recO | 1 | 649 | rv3863:rv3863 | 1 |
| 615 | rv2380c:mbtE | 1 | 650 | rv6034:tRNA-Gly32 | 1 |
| 616 | rv2154c:ftsW | 1 | 651 | rv1633:uvrB | 1 |
| 617 | rv2507:rv2507 | 1 | 652 | rv3531c:rv3531c | 1 |
| 618 | rv2699c:rv2699c | 1 | 653 | rv0544c:rv0544c | 1 |
| 619 | rv2173:idsA2 | 1 | 654 | rv0879c:rv0879c | 1 |
| 620 | rv6036:tRNA-Val34 | 1 | 655 | rv3207c:rv3207c | 1 |
| 621 | rv2594c:ruvC | 1 | 656 | rv2188c:pimB | 1 |
| 622 | rv3253c:rv3253c | 1 | 657 | rv3715c:recR | 1 |
| 623 | rv2520c:rv2520c | 1 | 658 | rv0555:menD | 1 |
| 624 | rv1831:rv1831 | 1 | 659 | rv3644c:rv3644c | 1 |
| 625 | rv3031:rv3031 | 1 | 660 | rv1631:coaE | 1 |
| 626 | rv1837c:glcB | 1 | 661 | rv1832:gcvB | 1 |
| 627 | rv0495c:rv0495c | 1 | 662 | rv0411c:glnH | 1 |
| 628 | rv0453:PPE11 | 1 | 663 | rv2726c:dapF | 1 |
| 629 | rv2206:rv2206 | 1 | 664 | rv1416:ribH | 1 |
| 630 | rv3754:tyrA | 1 | 665 | rv1657:argR | 1 |
| 631 | rv1712:cmk | 1 | 666 | rv2235:rv2235 | 1 |
| 632 | rv3510c:rv3510c | 1 | 667 | rv1658:argG | 1 |
| 633 | rv0849:rv0849 | 1 | 668 | rv0101:nrp | 1 |
| 634 | rv0542c:menE | 1 | 669 | rv3468c:rv3468c | 1 |

| | | | | | |
|---|---|---|---|---|---|
| 670 | rv3628:ppa | 1 | 705 | rv0382c:pyrE | 1 |
| 671 | rv3597c:lsr2 | 1 | 706 | rv0548c:menB | 1 |
| 672 | rv2257c:rv2257c | 1 | 707 | rv3789:rv3789 | 1 |
| 673 | rv3442c:rpsI | 1 | 708 | rv2139:pyrD | 1 |
| 674 | rv0467:icl1 | 1 | 709 | rv0220:lipC | 1 |
| 675 | rv0579:rv0579 | 1 | 710 | rv2777c:rv2777c | 1 |
| 676 | rv1619:rv1619 | 1 | 711 | rv3823c:mmpL8 | 1 |
| 677 | rv2172c:rv2172c | 1 | 712 | rv2747:argA | 1 |
| 678 | rv2438A:rv2438A | 1 | 713 | rv0057:rv0057 | 1 |
| 679 | rv2051c:ppm1 | 1 | 714 | rv0289:espG3 | 1 |
| 680 | rv1605:hisF | 1 | 715 | rv0723:rplO | 1 |
| 681 | rv0902c:prrB | 1 | 716 | rv2883c:pyrH | 1 |
| 682 | rv1005c:pabB | 1 | 717 | rv2842c:rv2842c | 1 |
| 683 | rv0013:trpG | 1 | 718 | rv2979c:rv2979c | 1 |
| 684 | rv2374c:hrcA | 1 | 719 | rv6003:tRNA-Gly4 | 1 |
| 685 | rv2097c:pafA | 1 | 720 | rv0796:rv0796 | 1 |
| 686 | rv1614:lgt | 1 | 721 | rv0704:rplB | 1 |
| 687 | rv3379c:dxs2 | 1 | 722 | rv3780:rv3780 | 1 |
| 688 | rv0188:rv0188 | 1 | 723 | rv3210c:rv3210c | 1 |
| 689 | rv0228:rv0228 | 1 | 724 | rv0164:TB18.5 | 1 |
| 690 | rv0780:purC | 1 | 725 | rv6014:tRNA-Ala15 | 1 |
| 691 | rv1409:ribG | 1 | 726 | rv1299:prfA | 1 |
| 692 | rv2977c:thiL | 1 | 727 | rv2993c:rv2993c | 1 |
| 693 | rv1371:rv1371 | 1 | 728 | rv3137:rv3137 | 1 |
| 694 | rv0416:thiS | 1 | 729 | rv3838c:pheA | 1 |
| 695 | rv2438c:nadE | 1 | 730 | rv3102c:ftsE | 1 |
| 696 | rv3713:cobQ2 | 1 | 731 | rv0809:purM | 1 |
| 697 | rv1659:argH | 1 | 732 | rv1240:mdh | 1 |
| 698 | rv2554c:rv2554c | 1 | 733 | rv1411c:lprG | 1 |
| 699 | rv2995c:leuB | 1 | 734 | rv0510:hemC | 1 |
| 700 | rv0102:rv0102 | 1 | 735 | rv1401:rv1401 | 1 |
| 701 | rv1202:dapE | 1 | 736 | rv2978c:rv2978c | 1 |
| 702 | rv2447c:folC | 1 | 737 | rv1600:hisC1 | 1 |
| 703 | rv2834c:ugpE | 1 | 738 | rv2664:rv2664 | 1 |
| 704 | rv1254:rv1254 | 1 | 739 | rv0803:purL | 1 |

| | | |
|---|---|---|
| 740 | rv3011c:gatA | 1 |
| 741 | rv6015:tRNA-Gln16 | 1 |
| 742 | rv0951:sucC | 1 |
| 743 | rv0998:rv0998 | 1 |
| 744 | rv1383:carA | 1 |
| 745 | rv2844:rv2844 | 1 |
| 746 | rv1795:eccD5 | 1 |
| 747 | rv0525:rv0525 | 1 |
| 748 | rv2746c:pgsA3 | 1 |
| 749 | rv2541:rv2541 | 1 |
| 750 | rv0423c:thiC | 1 |
| 751 | rv1559:ilvA | 1 |
| 752 | rv3041c:rv3041c | 1 |
| 753 | rv3058c:rv3058c | 1 |
| 754 | rv1173:fbiC | 1 |
| 755 | rv1465:rv1465 | 1 |
| 756 | rv0876c:rv0876c | 1 |
| 757 | rv0949:uvrD1 | 1 |
| 758 | rv3921c:rv3921c | 1 |
| 759 | rv1248c:rv1248c | 1 |
| 760 | rv2764c:thyA | 1 |
| 761 | rv3255c:manA | 1 |
| 762 | rv0505c:serB1 | 1 |
| 763 | rv0531:rv0531 | 1 |
| 764 | rv2199c:rv2199c | 1 |
| 765 | rv0479c:rv0479c | 1 |
| 766 | rv2697c:dut | 1 |
| 767 | rv3042c:serB2 | 1 |
| 768 | rv0352:dnaJ1 | 1 |
| 769 | rv1301:rv1301 | 1 |
| 770 | rv1384:carB | 1 |
| 771 | rv1093:glyA1 | 1 |
| 772 | rv2115c:mpa | 1 |
| 773 | rv0720:rplR | 1 |
| 774 | rv1464:csd | 1 |
| 775 | rv2465c:rpiB | 1 |
| 776 | rv3266c:rmlD | 1 |
| 777 | rv1836c:rv1836c | 1 |
| 778 | rv3378c:rv3378c | 1 |
| 779 | rv6022:tRNA-Leu20 | 1 |
| 780 | rv3907c:pcnA | 1 |
| 781 | rv1595:nadB | 1 |
| 782 | rv1822:pgsA2 | 1 |
| 783 | rv2192c:trpD | 1 |
| 784 | rv1229c:mrp | 1 |

**Table S4 – Primers used for KO construction and confirmation of KO and complementation**

| | |
|---|---|
| Rv0263c KO FP_ctgtgggatgtcgaccgac | Rv0263c KO RP_cagtcggtctcggtgaagtc |
| Rv0743c KO FP_gcggtcatgcacgtattgac | Rv0743c KO RP_ctggcccaacatcaacgaac |
| Rv0805 KO FP_cccctggtgatcgtggaac | Rv0805 KO RP_cttctgcctcaacgacggta |
| Rv0916c KO FP_gcaacgccagacaacacaaa | Rv0916c KO RP_caccgatccatacgagaccg |
| Rv1135A KO FP_ttgaccccagattccatggc | Rv1135A KO RP_gtctccaaccacgatgctga |
| Rv1339 KO FP_caaattggccgcacgattcc | Rv1339 KO RP_attcgatgagcaccgatccc |
| Rv2171 KO FP_tgggcttcaccgacatcatc | Rv2171 KO RP_cgacttgtacaagcgtgtgc |
| Rv2287 KO FP_ctcgatgacattgtgccgtg | Rv2287 KO RP_ccagtgcaatgaaacgctga |
| Rv2515c KO FP_ttaatcggctggtcgggaac | Rv2515c KO RP_cctcatcgaccgacctgatc |
| Rv2934 KO FP_atggatttgctcgccgatct | Rv2934 KO RP_gaacgccgaaacatccttgg |
| Rv3484 KO FP_tggccccatactagacgtca | Rv3484 KO RP_ggacaggtcatcaacgtcga |
| Rv3842c KO FP_tgaacacagacgggcgttc | Rv3842c KO RP_tgcagtggttcttgcaggaa |
| Rv3896c KO FP_gatccttgttgtgccgaacg | Rv3896c KO RP_acgttgagcagcatcatcca |
| Rv1129c KO FP_tcttgtaggccatgtgctcg | Rv1129c KO RP_atctcctcccgagtggaaca |
| Rv1131 KO FP_tggtgtagtggaacccgttg | Rv1131 KO RP_gcgacgtaaaacaggtgcag |
| Rv1427c KO FP_ccgatcaacctcaccaagca | Rv1427c KO RP_ggtacactcctggcacttcc |
| Rv1888c KO FP_attactgcacgtcgatccgg | Rv1888c KO RP_ctagagcacatacccggcg |
| Rv2002 KO FP_cattcgttacgagcagcacg | Rv2002 KO RP_cccactcgagtcggaatagc |
| Rv2103c KO FP_tcaagaaggccctcaacgac | Rv2103c KO RP_ctagaccgctattacgcccg |
| Rv2144c KO FP_caaccggggcaaaaactagc | Rv2144c KO RP_tctcgtgaggtgactgggat |
| Rv2482c KO FP_acgaaggcggctacaaagaa | Rv2482c KO RP_cccactgacccagttcactg |
| Rv2517c KO FP_cactcactggctacatcgct | Rv2517c KO RP_cgagataccagtgcgtcgaa |
| Rv2953 KO FP_cacagaagtcacgggacctg | Rv2953 KO RP_gtgggttgtcgtcaggagtt |
| Rv2970c KO FP_aaagacatccagcctgacgg | Rv2970c KO RP_gctcgatttgtcggcttgtc |
| Rv3156 KO FP_atcgcgttcttcaccatggt | Rv3156 KO RP_cgggtggtagcaggatgatc |
| Rv3268 KO FP_gtagtgatccgaacggcgtc | Rv3268 KO RP_tcgccattgcctgtaccttt |
| Rv3370c KO FP_ttcgcaggtgaagatgtgct | Rv3370c KO RP_tgaaatccacgacgcgtttg |
| Rv3479 KO FP_cattcgaaaagccgtcaccg | Rv3479 KO RP_agccaggctctcaacgttac |
| Rv3741c KO FP_ctggtggagatcgaacgctt | Rv3741c KO RP_cctgataggtttcccgcaca |
| Rv3779 KO FP_ccatgccagccaccttagaa | Rv3779 KO RP_tgcttgatcatggtgccgat |
| Rv0263c Comp FP_caccaacagcattcaccacg | Rv0263c Comp RP_atgtcctcgtcggtgatgac |
| Rv0743c Comp FP_atgtggacgtgctggtactc | Rv0743c Comp RP_caaaacccgagcctggatca |
| Rv0805 Comp FP_gggccgaactacgcaaattc | Rv0805 Comp RP_cgatgaaaatgccgctctcg |
| Rv0916c Comp FP_cgtgagtgctcggaacacag | Rv0916c Comp RP_actcggtagtgcttggtgtg |
| Rv1135A Comp FP_cttaccccctgttcaacccc | Rv1135A Comp RP_atcgtctgtaagccgtagcg |
| Rv1339 Comp FP_ttggttatcgacttcggcgg | Rv1339 Comp RP_acaggtgtagatcgggtgga |
| Rv2171 Comp FP_cgagcctcgatcaccatctc | Rv2171 Comp RP_gatgcccagccagatgcc |

| | |
|---|---|
| Rv2287 Comp FP_cattgctacgccgcagaatc | Rv2287 Comp RP_acggtgaccagaatgacgac |
| Rv2515c Comp FP_caaggccttgatcgaggtca | Rv2515c Comp RP_cgattccaagcgtgcttagc |
| Rv2934 Comp FP_cggccgatttttatgccctg | Rv2934 Comp RP_gagtaccagccaacttcccc |
| Rv3484 Comp FP_cacgctgatacttgtgcacg | Rv3484 Comp RP_aaggtgcccgaatcctgaag |
| Rv3842c Comp FP_gcatctggtctgtgtgcatg | Rv3842c Comp RP_tcgacgttccagcagtacac |
| Rv3896c Comp FP_gatatcctgggtggccaacg | Rv3896c Comp RP_ggcgttgtagttcgattcgc |
| Rv1129c Comp FP_gcgcagtatttctcctcgga | Rv1129c Comp RP_cgagatcaggtctgactggc |
| Rv1131 Comp FP_gacatcaaaaagggcctcgc | Rv1131 Comp RP_gctcggcgtagaggatcatc |
| Rv1427c Comp FP_ttcatggaccaattcggcga | Rv1427c Comp RP_ttgtctcggacatgctgctt |
| Rv1888c Comp FP_tggcagtggctagtcaagtg | Rv1888c Comp RP_gcaggatcaccagtagcgaa |
| Rv2002 Comp FP_aaggcgcaaaggttgtgttc | Rv2002 Comp RP_ccggtggaatagctcgactc |
| Rv2103c Comp FP_tgagcaccacaagccgtc | Rv2103c Comp RP_gtaggacacgatgctggcg |
| Rv2144c Comp FP_cagctagtggcctgggtatg | Rv2144c Comp RP_ctcctcactgccgatgacac |
| Rv2482c Comp FP_tcgatcagctgcacgagatc | Rv2482c Comp RP_cgtcaggcgctatgtaccat |
| Rv2517c Comp FP_gacggttgaggatgacgcc | Rv2517c Comp RP_tcactgtgcgccttccttc |
| Rv2953 Comp FP_gcggattcgattcgatccct | Rv2953 Comp RP_taatcgtcggggtagtcggt |
| Rv2970c Comp FP_gacgtgaccgacctgtcaat | Rv2970c Comp RP_cgacatcggaatccctgagg |
| Rv3156 Comp FP_ttctggtaccacaagccgtc | Rv3156 Comp RP_agatgcatgatcgcaaacgc |
| Rv3268 Comp FP_ggtgacactggctaactggg | Rv3268 Comp RP_gtccaccaactcatcgggtc |
| Rv3370c Comp FP_gttctactcggcgtggttca | Rv3370c Comp RP_aggaactgggtcggatagct |

**Table S5**

| **Gene** | **Stgae 1** | **Stage 2** | **Stage 3** | **Stgae 4** |
|---|---|---|---|---|
| kstR1: *rv3574* | 0 | 0 | 1 | 0 |

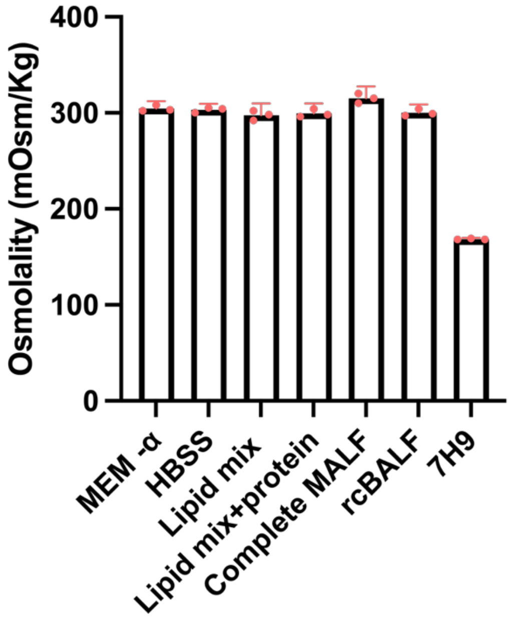


**Figure S1: Osmolality of MALF, rcBALF, 7H9, and MALF components.** Osmolality (mOsm/kg) was measured for various media and fluid formulations. Complete MALF and its individual components exhibited physiological osmolality (~300 mOsm/kg), comparable to rcBALF. Complete 7H9 medium showed a non-physiologic osmolality of approximately 160 mOsm/kg.

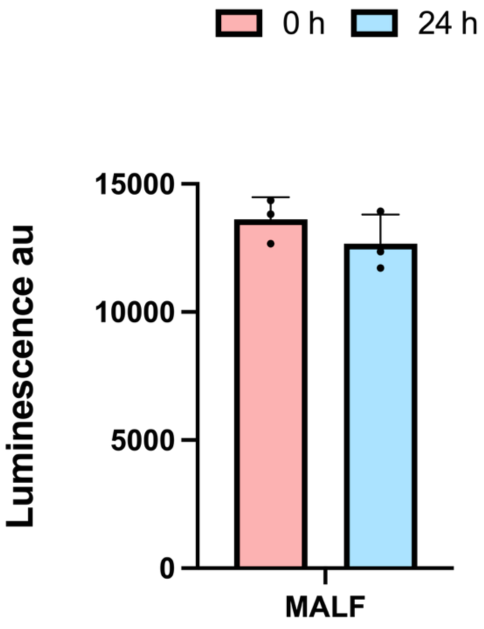


**Figure S2. Non-cytotoxicity of MALF to human lung cells.** BALF collected on two different days from different donors was centrifuged. The pelleted cells were put into culture with (describe the medium) for 3 days. The medium was replaced with MALF and cytotoxicity was measured 24 h later as described in Methods. Results are pooled means ± SEM for triplicates in the two independent experiments.

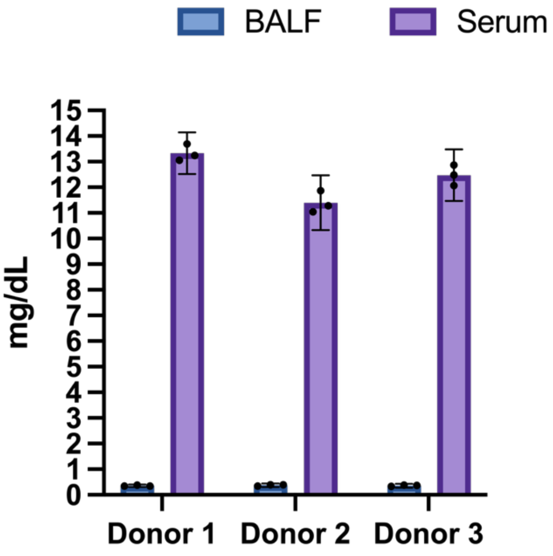


**Figure S3. BUN concentrations in matched serum and BALF**. Comparison of blood urea nitrogen (BUN) levels (mg/dL) in serum (blue) and BALF (purple) from three healthy donors, reflecting the degree to which the lavage diluted the ALF, where the concentration of urea should match that in the donor’s serum.

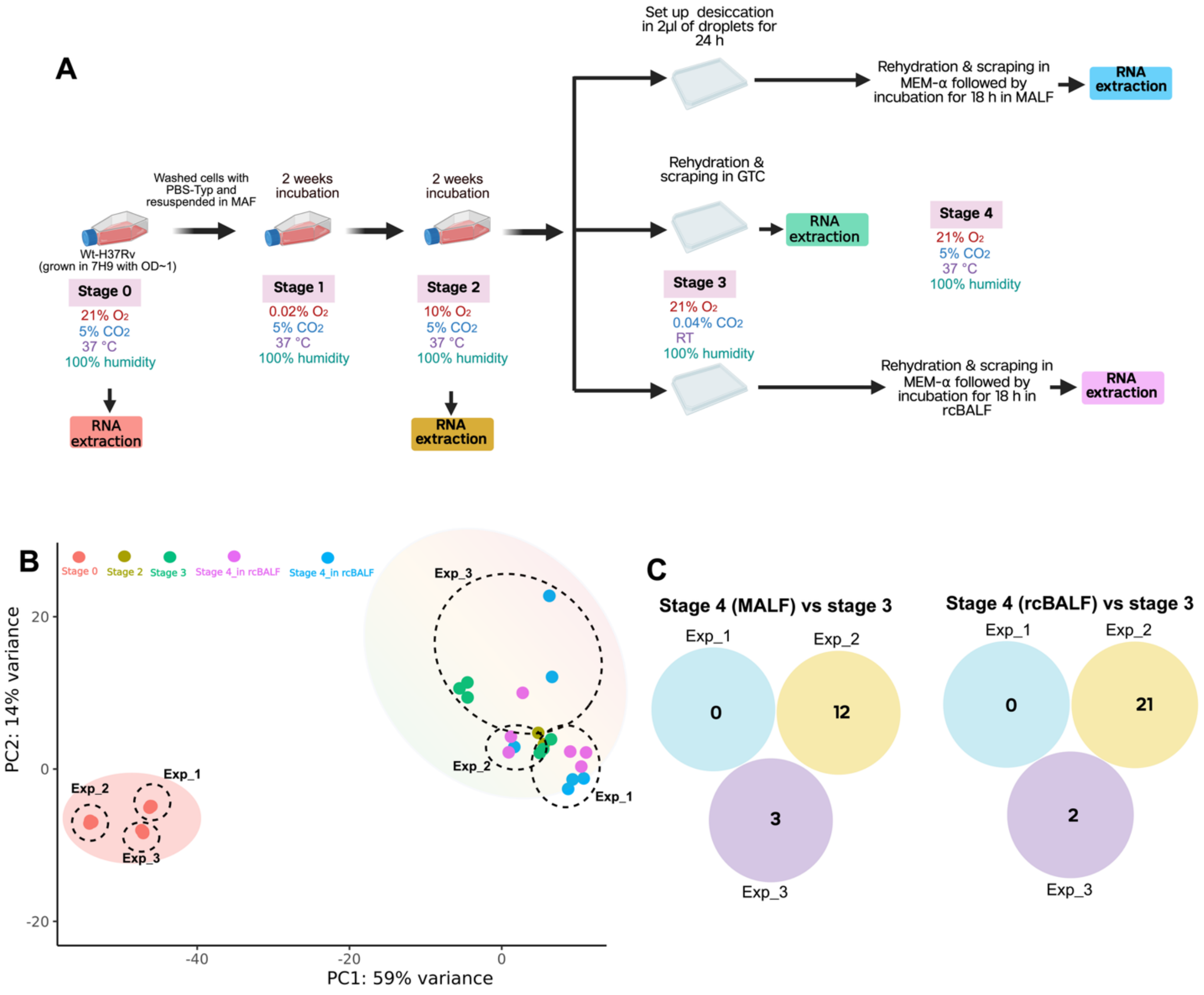


**Figure S4. Transcriptome of Mtb in the transmission model. (A)** Schematic of the experimental setup for the RNA experiment. **(B)** PCA plot comparing transcriptomes of Mtb from log-phase culture in 7H9 (stage 0) and Mtb after 2 weeks of incubation in MAF in 0.2% $O_2$, followed by an additional 2 weeks in MAF in 10% $O_2$ (stage 2) and desiccation air as recovered in GTC after desiccation (stage 3) or as rehydrated briefly in MEMα and then incubated for 18 h in rcBALF (stage 4) or MALF (stage 4). **(C)** Venn diagrams representing the number of significantly regulated genes when comparing stage 4 (MALF or rcBALF) against stage 3. Significance for these differentially expressed genes (DEGs) is defined by a padj value ≤ 0.05, log2FC ≥ 1.5 or log2FC ≤ -1.5. Data represents results from three independent RNA-seq experiments (Exp_1, Exp_2, and Exp_3)

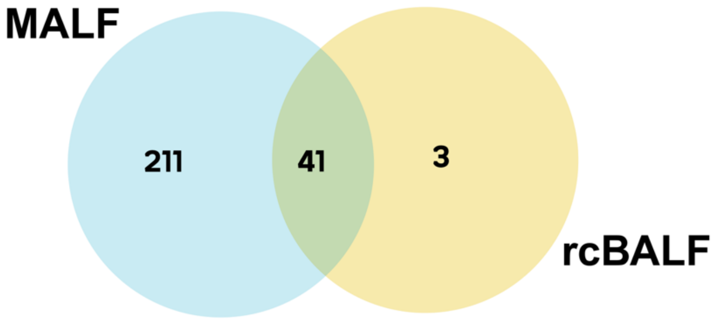


**Figure S5. Comparison of differentially expressed genes (DEGs) in Mtb incubated in MALF or rcBALF.** Venn diagram illustrating the overlap of DEGs identified via RNA-seq analysis. The blue circle represents genes consistently identified across three samples for MALF, while the yellow circle represents genes consistently identified from rcBALF. The central intersection indicates 41 common genes that are consistently and significantly differentially expressed in Mtb after incubating in both fluids**.** (padj value ≤ 0.05, log2FC ≥ 1.5 or log2FC ≤ -1.5).

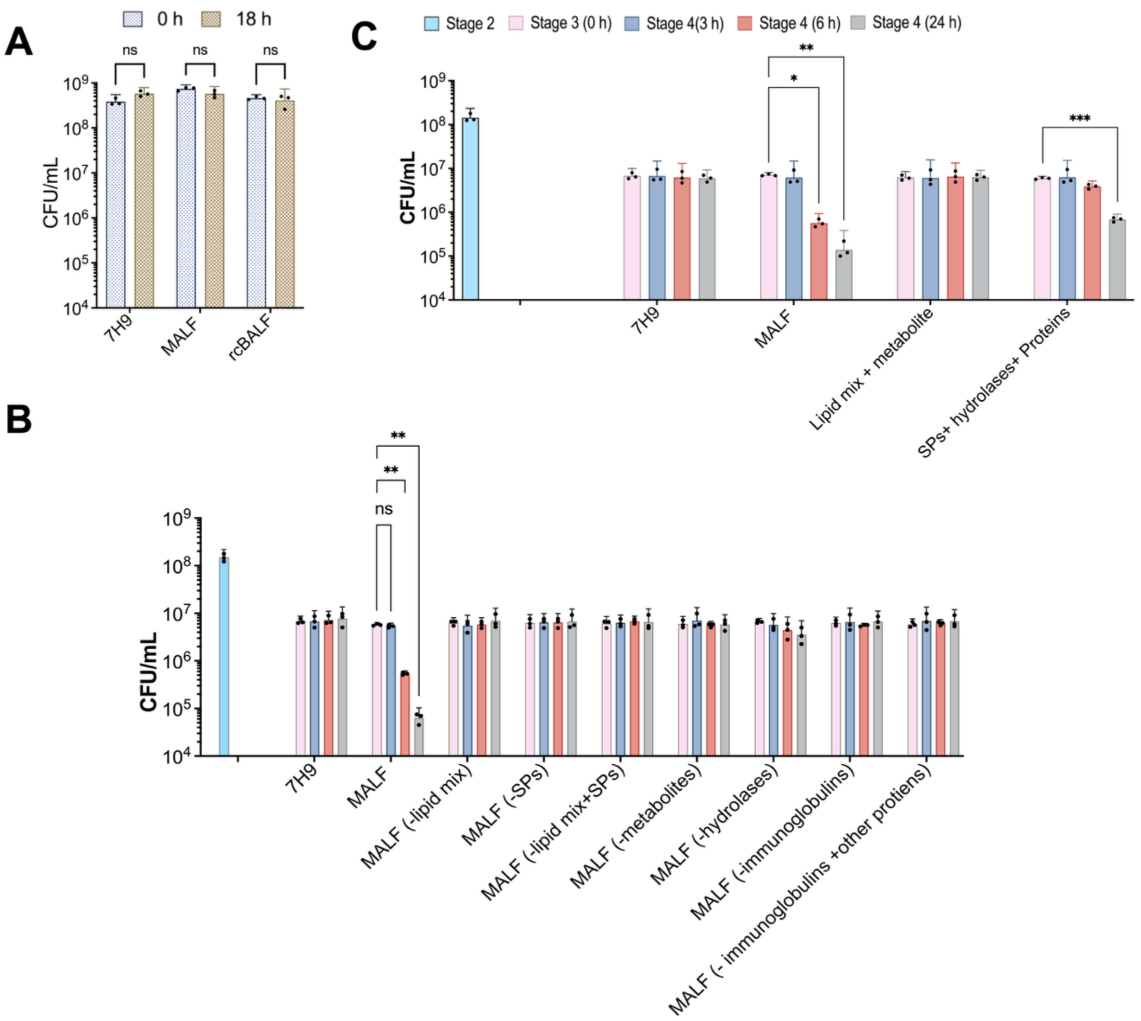


**Figure S6. Effect of MALF & rcBALF on survival of log-phase and transmission-modeled Mtb.** (**A**) CFU/mL of stage 0 incubated Mtb remained stable after 18h in 7H9, MALF, and rcBALF under 21% $O_2$ 5% $CO_2$. (**B, C**) Survival of desiccated Mtb rehydrated in various MALF formulations or 7H9 at 3, 6, and 24 h. The key above panel C applies also to panel B. Data represent mean + SEM from three biological repeats. Statistical significance was determined by two-way ANOVA followed by Tukey'smultiple comparisons test (*p < 0.05, **p < 0.01, ***p < 0.001).

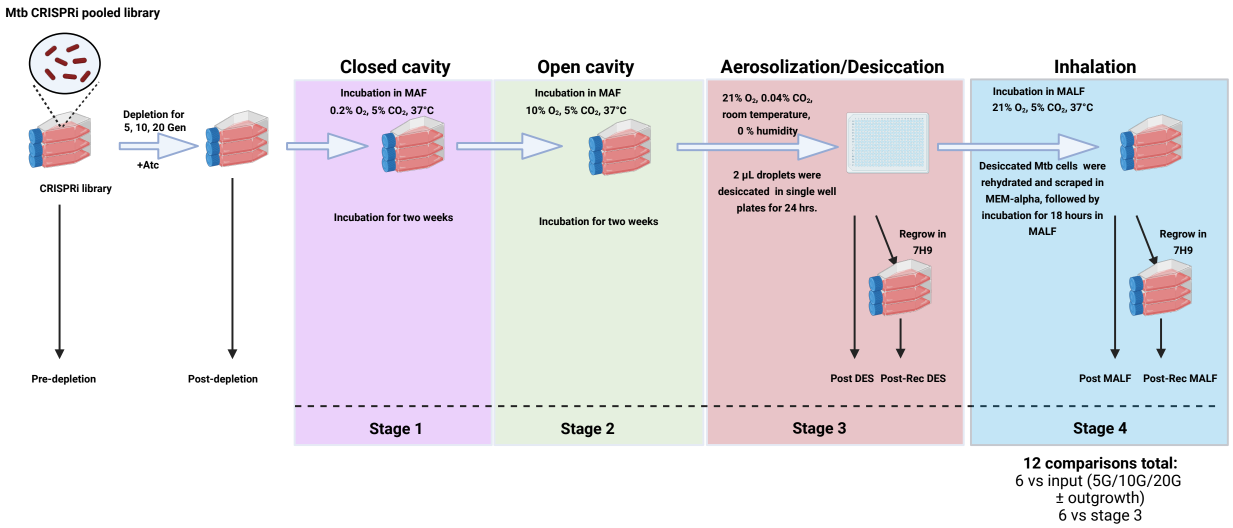


**Figure S7**. **Genome-wide CRISPRi screening across the aerosolization-to-inhalation stages**. Schematic of the CRISPRi screen. A pooled library was passaged through the first two stages, followed by desiccation (stage 3) and incubation in MALF (stage 4). Arrows indicate the stages at which samples were collected for genomic DNA isolation.

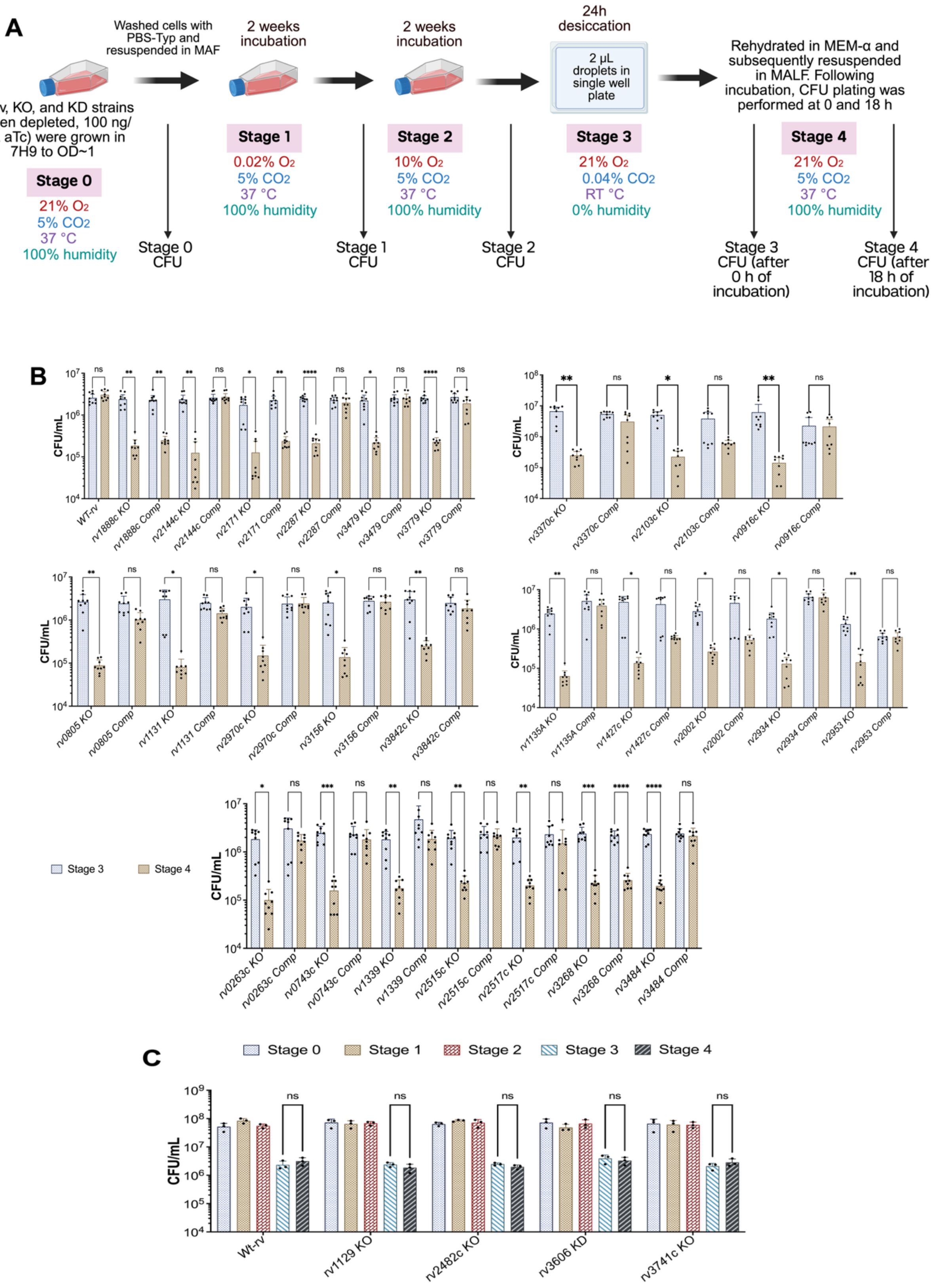
A
Washed cells with PBS-Typ and resuspended in MAF
2 weeks incubation
2 weeks incubation
24h desiccation
2 µL droplets in single well plate
Rehydrated in MEM-α and subsequently resuspended in MALF. Following incubation, CFU plating was performed at 0 and 18 h
Wt-rv, KO, and KD strains (5-gen depleted, 100 ng/mL aTc) were grown in 7H9 to OD~1
Stage 0
21% O2
5% CO2
37 °C
100% humidity
Stage 0 CFU
Stage 1
0.02% O2
5% CO2
37 °C
100% humidity
Stage 1 CFU
Stage 2
10% O2
5% CO2
37 °C
100% humidity
Stage 2 CFU
Stage 3
21% O2
0.04% CO2
RT °C
0% humidity
Stage 3 CFU (after 0 h of incubation)
Stage 4
21% O2
5% CO2
37 °C
100% humidity
Stage 4 CFU (after 18 h of incubation)
B
CFU/mL
WT-rv
rv1888c KO
rv1888c Comp
rv2144c KO
rv2144c Comp
rv2171 KO
rv2171 Comp
rv2287 KO
rv2287 Comp
rv3479 KO
rv3479 Comp
rv3779 KO
rv3779 Comp
rv3370c KO
rv3370c Comp
rv2103c KO
rv2103c Comp
rv0916c KO
rv0916c Comp
rv0805 KO
rv0805 Comp
rv1131 KO
rv1131 Comp
rv2970c KO
rv2970c Comp
rv3156 KO
rv3156 Comp
rv3842c KO
rv3842c Comp
rv1135A KO
rv1135A Comp
rv1427c KO
rv1427c Comp
rv2002 KO
rv2002 Comp
rv2934 KO
rv2934 Comp
rv2953 KO
rv2953 Comp
Stage 3
Stage 4
rv0263c KO
rv0263c Comp
rv0743c KO
rv0743c Comp
rv1339 KO
rv1339 Comp
rv2515c KO
rv2515c Comp
rv2517c KO
rv2517c Comp
rv3268 KO
rv3268 Comp
rv3484 KO
rv3484 Comp
C
Stage 0
Stage 1
Stage 2
Stage 3
Stage 4
CFU/mL
Wt-rv
rv1129 KO
rv2482c KO
rv3606 KD
rv3741c KO

**Figure S8. Experimental workflow and Mtb survival during different stages of transmission model.** (**A**) Schematic of the experimental design. Starting from log phase cultures in stage 0 (7H9 under 21% $O_2$, 5% $CO_2$), knockout strains and CRISPRi strains that were depleted for five generations were washed, resuspended in MAF and transitioned through sequential 2-week incubations in 0.2% $O_2$, 5% $CO_2$ (stage 1) followed by 10% $O_2$, 5% $CO_2$ (stage 2). Samples were then desiccated for 24 hours, after which they were rehydrated in MALF under 21% $O_2$, 5% $CO_2$ for CFU plating at 0 (stage 3) and 18 (stage 4) hours. **(B)** CFU of wild type Mtb H37Rv, KO & complemented strains at different stages. **(C**) CFU for wild type Mtb H37Rv, knockout strains and the knockdown strains for *rv1129, rv2482c, rv3606* and *rv3741c* at different stages. Statistical significance was determined by two-way ANOVA followed by Tukey'smultiple comparisons test (*p < 0.05, **p < 0.01, ***p < 0.001).

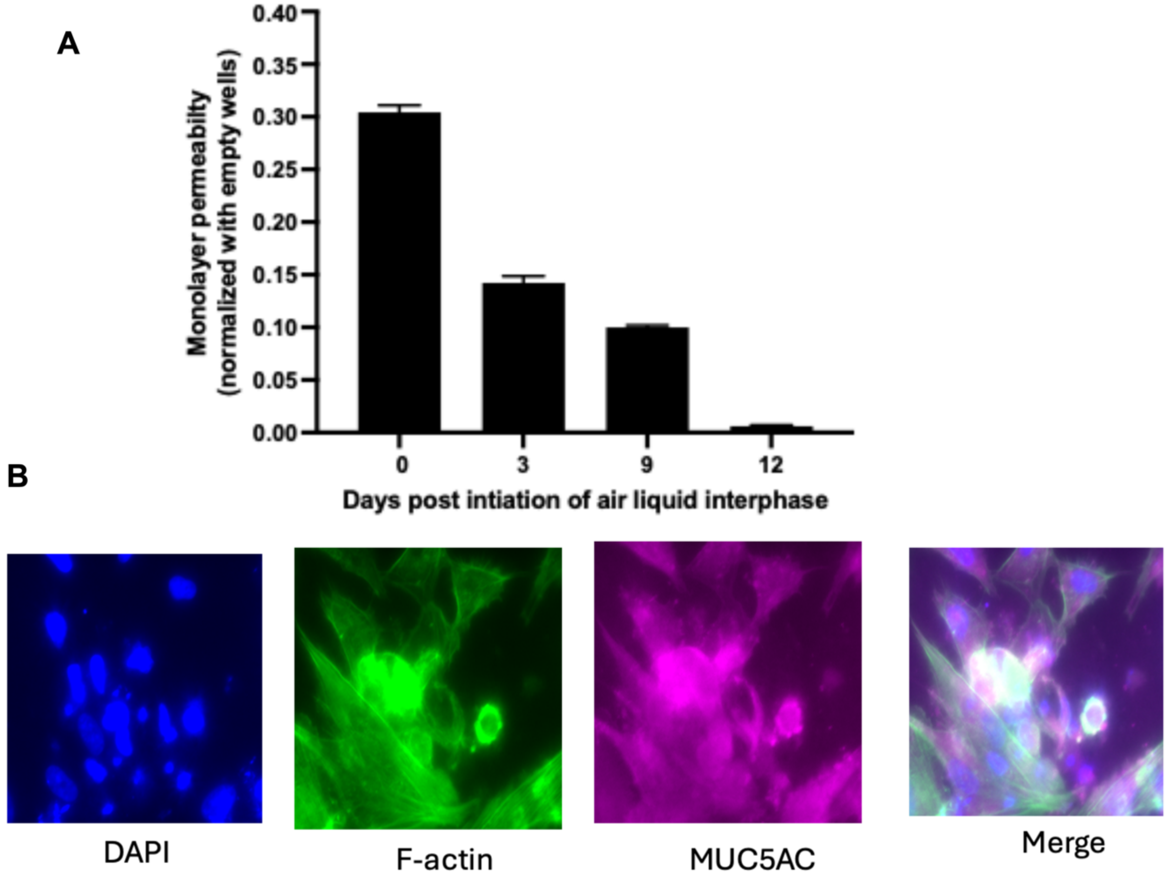


**Figure S9. Characterization of NuLi-1 Lung epithelial cell differentiation in air-liquid Interface (ALI) culture. (A)** Fluorescein permeability assay showing the development of barrier function over a 12-day period. The progressive decrease in permeability (normalized to cell-free wells) indicates the formation of a confluent, tight-junctionreinforced monolayer post-initiation of ALI. **(B)** Representative confocal microscopy images of NuLi-1 cells at day 12 of ALI. Fixed cultures were stained with DAPI (blue) to visualize nuclei, phalloidin for F-actin (green) to show cytoskeletal structure, and antibodies against MUC5AC (magenta) to identify differentiated, mucus-secreting goblet cells.

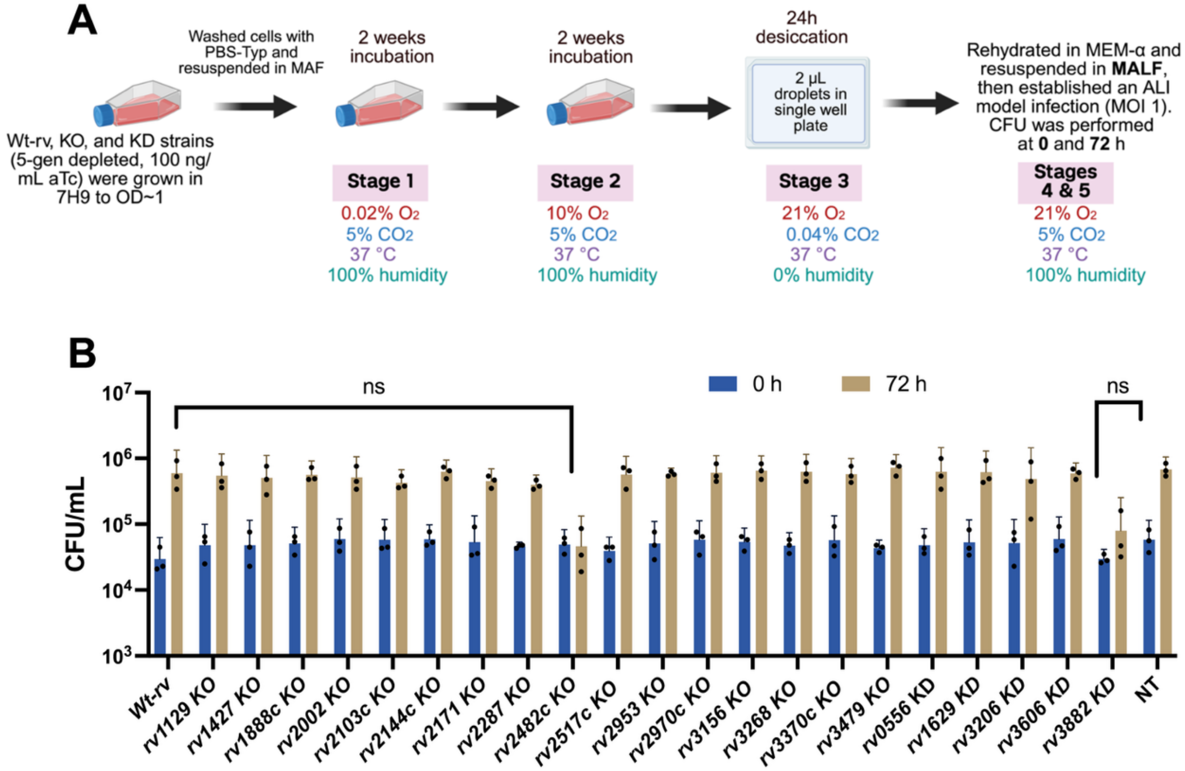


**Figure S10. Survival of Mtb strains following sequential stages of transmission in an ALI model infection. (A)** Schematic of the experimental workflow. Wt-rv, KO, and KD strains were grown to OD ~1, resuspended in MAF, and incubated through sequential stages of hypoxia (stages 1 & 2), 24 h desiccation (stage 3), and rehydration in MALF for infection of an ALI model (stages 4 & 5) **(B)** Bacterial recovery post-infection. CFU/mL was measured at 0 h and 72 h post-infection. Data are shown as mean ± SD from three biological replicates. Statistical significance was determined by two-way ANOVA followed by Sidak's multiple comparison test (ns, not significant).

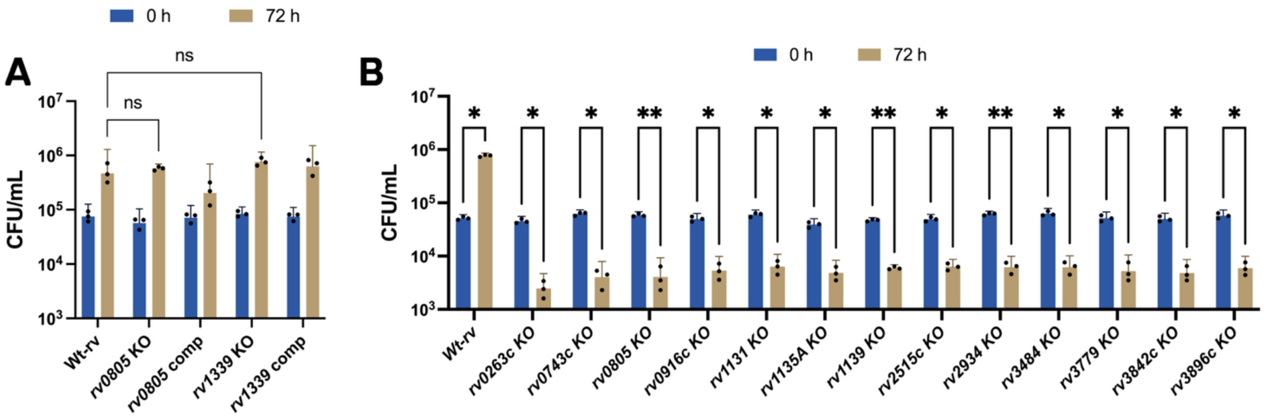


**Figure S11. Intracellular survival of various Mtb H37*rv* mutant strains in two different models of infection. (A)** Intracellular survival in a monolayer of alveolar macrophage-like cells. Mtb strains came from stage 0 culture; that is, they were not incubated in the sequential stages of the transmission model prior to infection. CFU/mL were measured at 0 hours (blue bars) and 72 hours (tan bars) post-infection with *Wt-rv*, KO and complemented strains.(**B).** Intracellular survival in an air-liquid interface (ALI)-based infection model using human monocyte-derived macrophages. *Mtb strains* were passaged through stages 1 to 4 of the transmission model before infection. CFU/mL were measured at 0 hours (blue bars) and 72 hours (tan bars) post-infection with *Wt-rv* or KO strains. Statistical significance was determined by two-way ANOVA followed by Sidak's multiple comparison test:(* $p < 0.05$, ** $p < 0.01$).

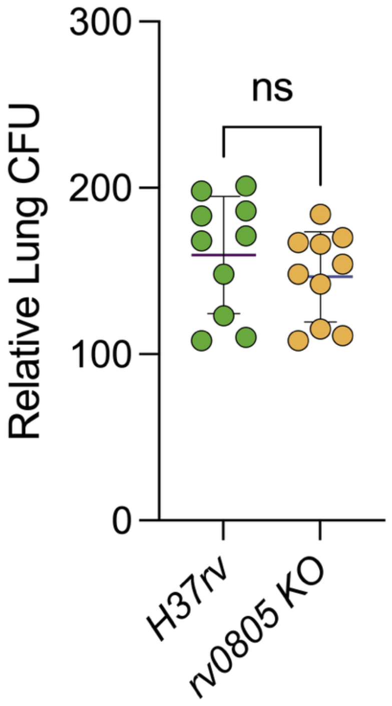


**Figure S12.** Lung CFU 24 h after infection in mice exposed to respirable-size aerosols in the TSS, using only Stage 0–incubated wild-type (green) and *rv0805* knockout (yellow) Mtb strains. Data were analyzed by two-tailed Welch's t test. $p$<0.05 was considered statistically significant.

| No. | Gene | Stage 1 | Stage 2 | Stage 3 | Stage 4 | Enzyme |
|---|---|---|---|---|---|---|
| 1 | rv0891c | 0 | 0 | 0 | 0 | ACs |
| 2 | rv1120c | 3 | 2 | 3 | 0 | |
| 3 | rv1264 | 1 | 0 | 0 | 0 | |
| 4 | rv1318c | 1 | 0 | 0 | 0 | |
| 5 | rv1319c | 0 | 0 | 0 | 0 | |
| 6 | rv1320c | 1 | 0 | 0 | 0 | |
| 7 | rv1358 | 1 | 2 | 0 | 0 | |
| 8 | rv1359 | 2 | 0 | 1 | 0 | |
| 9 | rv1625c | 1 | 0 | 1 | 0 | |
| 10 | rv1647 | 0 | 1 | 1 | 0 | |
| 11 | rv1900c | 0 | 0 | 0 | 0 | |
| 12 | rv2212 | 0 | 0 | 0 | 0 | |
| 13 | rv2435c | 2 | 0 | 0 | 0 | |
| 14 | rv2488c | 3 | 2 | 4 | 0 | |
| 15 | rv3645 | 6 | 6 | 11 | 6 | |
| 16 | rv0805 | 0 | 0 | 0 | 1 | PDEs |
| 17 | rv1339 | 4 | 1 | 0 | 1 | |

**Figure S13:** Adenyyl cyclases (Acs) and phosphodiesterases (PDEs) identified as potentially required for Mtb's survival at different stages of modeled transmission. Numerical values indicate NOSDs, representing the number of comparisons in which the sgRNA count for the indicated gene at the indicated stage was statistically significantly below its sgRNA count at one or more earlier stages.

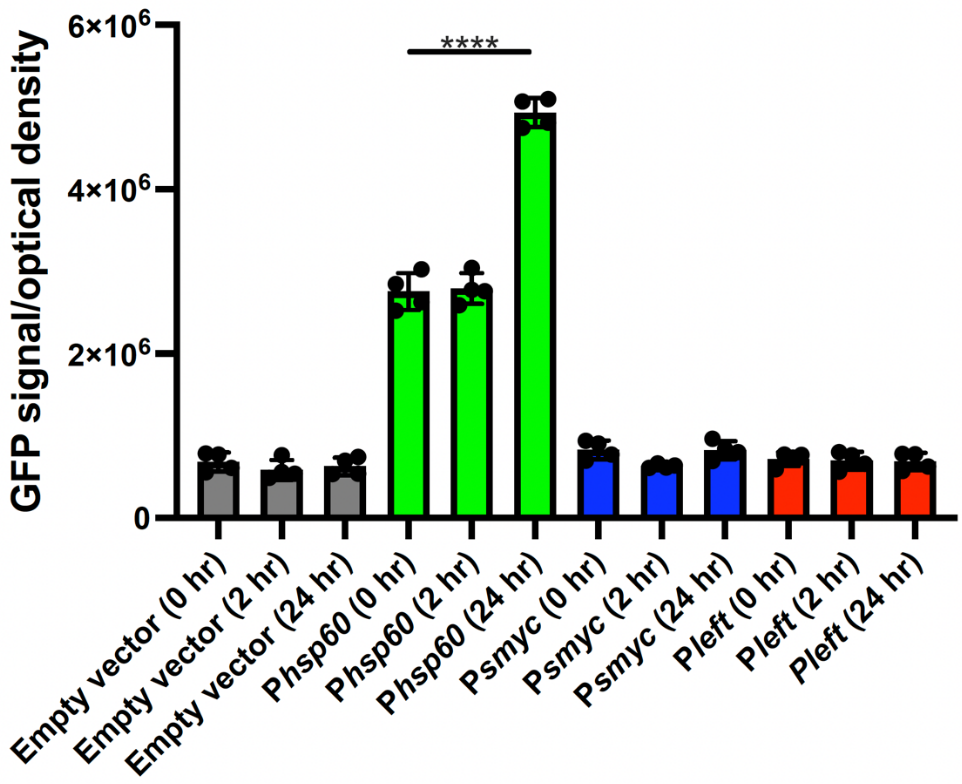


**Figure S14. Optimizing G-Flamp1 expression in Mtb and confirming that the G-Flamp1 signal increases when cAMP levels are modulated.** The DNA sequence encoding a GFP-based cAMP sensor (G-Flamp1) was codon optimized for expression in Mtb while maintaining the G-Flamp1 primary amino acid sequence [16]. We expressed the G-Flamp1 reporter in Mtb under control of different promoters (P*smyc*, P*hsp60*, and P*left*) and quantified the G-Flamp1 signal in the presence of the Rv1625/cya agonist (V59) to increase bacterial levels of cAMP [17]. The G-Flamp1 signals were quantified with a PerkinElmer Envision plate reader and normalized for bacterial numbers using absorbance measurement at 600 nm and reported as GFP signal/optical density. We chose to express the G-Flamp1 reporter from the P*hsp60* promoter. The reporter signal increased when the bacteria were treated with V59- [17] at 10 mM for 24 hours. Results are means ± SEM from 2 independent experiments. Significance is indicated by asterisks (****$p < 0.0001$). Statistical significance was determined by one-way Anova followed by Sidak's multiple comparison test.

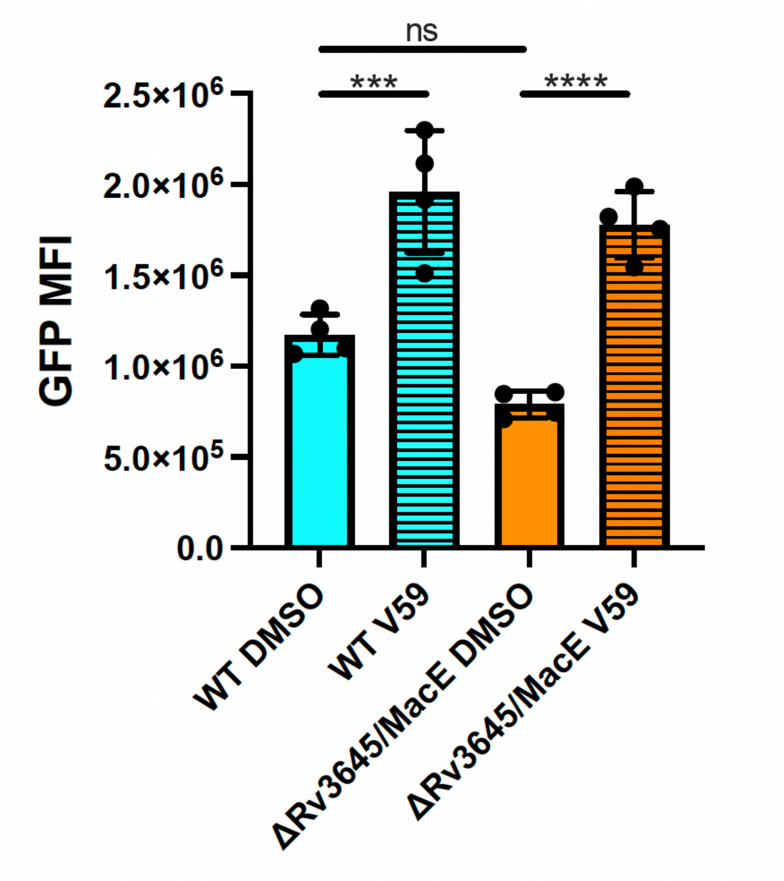


**Figure S15. Creating an mCherry enabled G-Flamp1 reporter.** The P*hsp60*::G-Flamp1 reporter plasmid was engineered to constitutively express mCherry for flow cytometric and image-based analysis. We introduced the mCherry/G-Flamp1 reporter into WT Mtb and an Mtb mutant lacking the adenylyl cyclase Rv3645/MacE, a strain that produces very little cAMP in standard *in vitro* growth conditions [18]. The G-Flamp1 signal was quantified from paraformaldehyde fixed cells using a BD Accuri flow cytometer. For this analysis, ~20,000 mCherry positive bacterial cells were detected in each sample followed by quantification of the mean fluorescent intensity in the GFP channel. Relative to WT, the G-Flamp1 signal in the Rv3645/MacE mutant was slightly decreased without statistical significance, indicating the bottom signal threshold for this reporter in Mtb. Following V59 treatment, both WT and the Rv3645/MacE mutant produced similar G-Flamp1 signal levels, indicating the upper signal threshold for this reporter in Mtb. Results are means ± SEM from 2 independent experiments.Significance is indicated by asterisks (***$p < 0.001$, ****$p < 0.0001$). Statistical significance was determined by one-way Anova followed by Sidak's multiple comparison test.

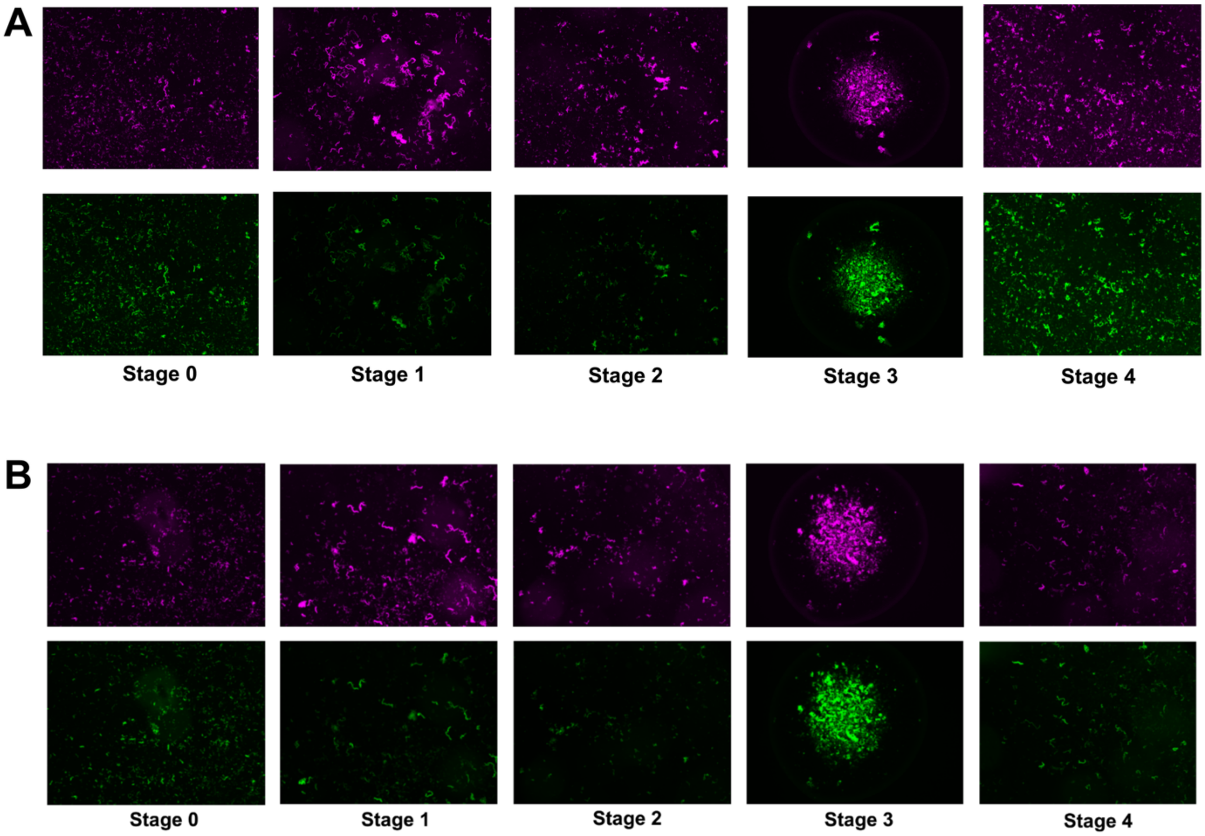


**Figure S16. cAMP levels in *rv0805* and *rv1339* Mtb mutants across transmission stages.** Fluorescence microscopy of **(A)** *rv0805* and **(B)** *rv1339* knockout strains during stages 0–4 of a transmission model. Magenta (top rows): constitutive mCherry expression serves as a control for total bacterial count. Green (bottom rows): GFP intensity reports high intracellular cAMP levels. Data are representative of three biological replicates.

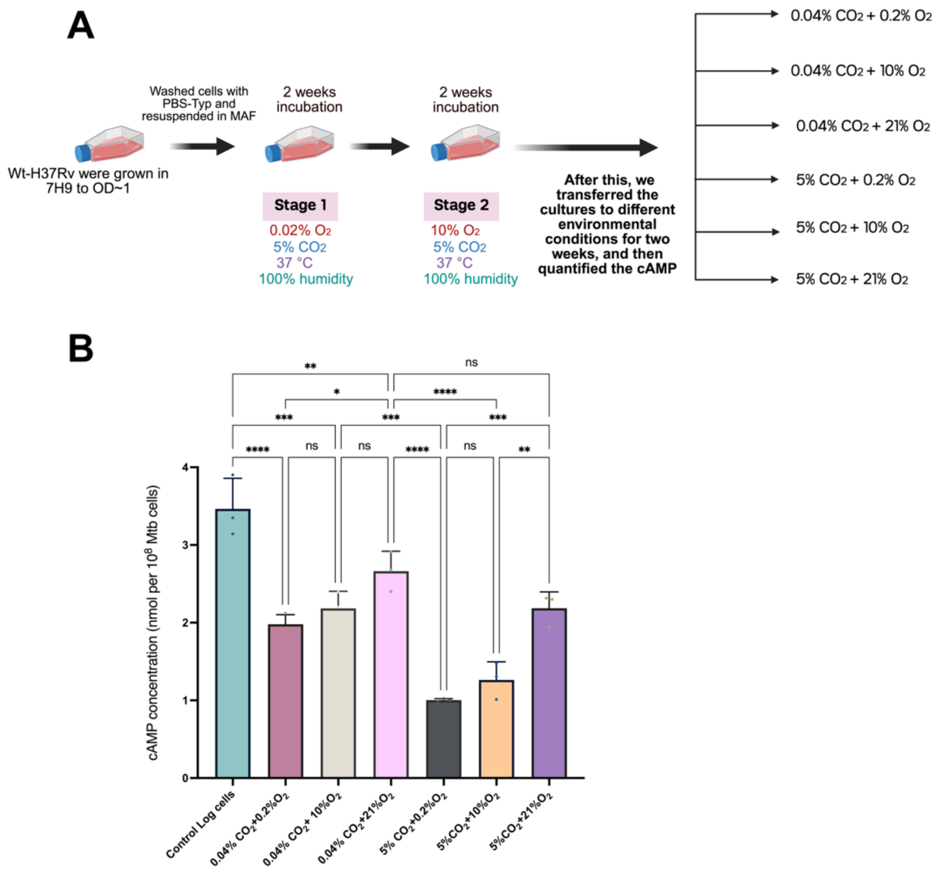


**Figure. S17. cAMP concentrations in Mtb in varying gaseous and liquid environments. (A)** Schematic. After Mtb in MAF was incubated for 2 weeks each in the atmospheres of stages 1 and 2, samples were incubated in the indicated atmospheres for another 2 weeks. **(B)** cAMP levels were then measured and compared to levels in Mtb at log phase in 7H9 under 21% $O_2$, 5% $CO_2$ (stage 0). Results are means mean ± SEM of three independent experiments. Statistical significance was determined by two-way ANOVA followed by Tukey'smultiple comparisons test (*$p < 0.05$, **$p < 0.01$, ***$p < 0.001$, ****$p < 0.0001$).

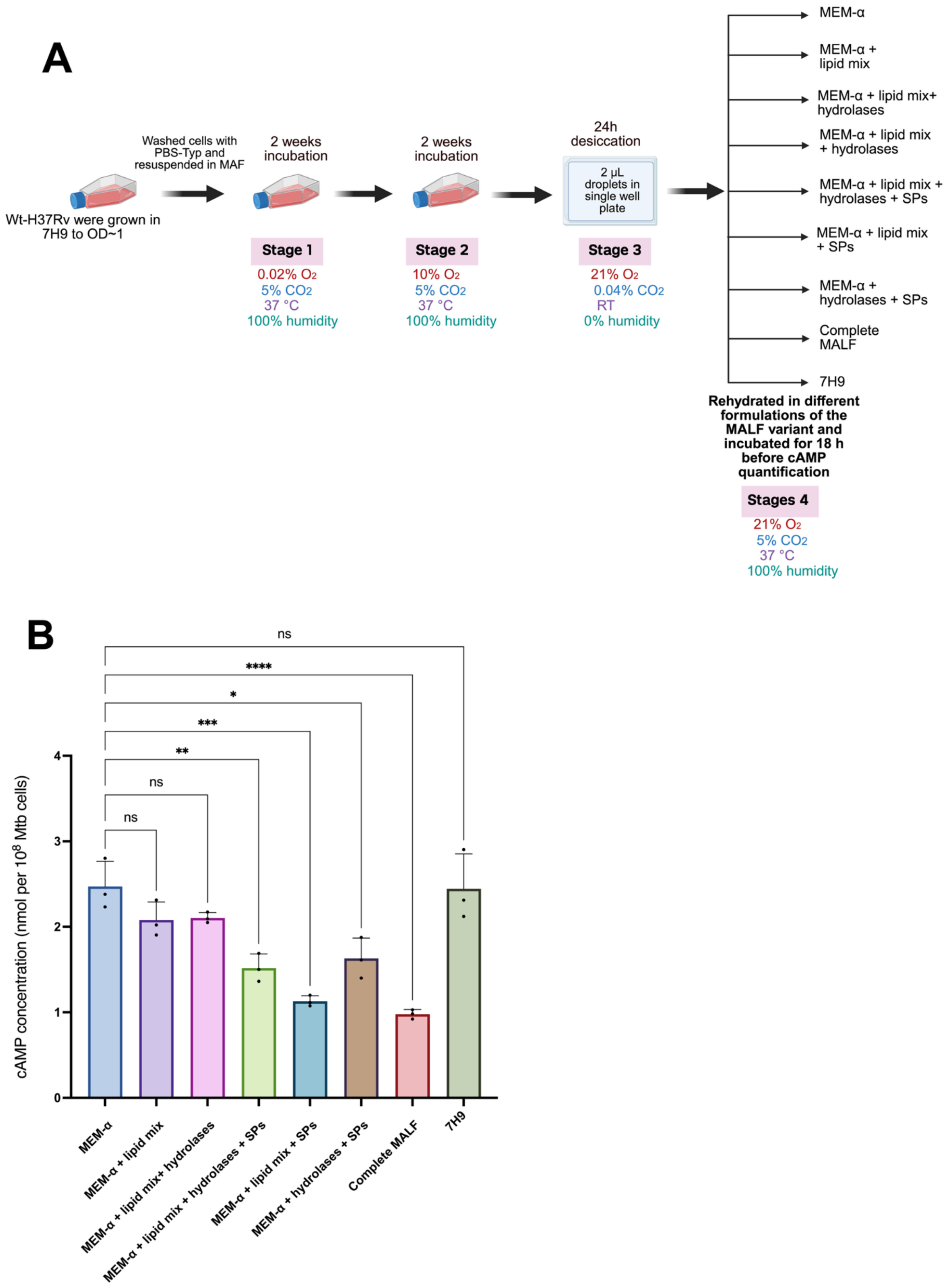
A
Wt-H37Rv were grown in 7H9 to OD~1
Washed cells with PBS-Typ and resuspended in MAF
2 weeks incubation
Stage 1
0.02% $O_2$
5% $CO_2$
37 °C
100% humidity
2 weeks incubation
Stage 2
10% $O_2$
5% $CO_2$
37 °C
100% humidity
24h desiccation
2 µL droplets in single well plate
Stage 3
21% $O_2$
0.04% $CO_2$
RT
0% humidity
MEM-α
MEM-α + lipid mix
MEM-α + lipid mix+ hydrolases
MEM-α + lipid mix + hydrolases
MEM-α + lipid mix + hydrolases + SPs
MEM-α + lipid mix + SPs
MEM-α + hydrolases + SPs
Complete MALF
7H9
Rehydrated in different formulations of the MALF variant and incubated for 18 h before cAMP quantification
Stages 4
21% $O_2$
5% $CO_2$
37 °C
100% humidity
B
cAMP concentration (nmol per $10^8$ Mtb cells)
ns
****
*
***
**
ns
ns
0
1
2
3
4
MEM-α
MEM-α + lipid mix
MEM-α + lipid mix+ hydrolases
MEM-α + lipid mix + hydrolases + SPs
MEM-α + lipid mix + SPs
MEM-α + hydrolases + SPs
Complete MALF
7H9

**Figure S18. Effect of MALF components on cAMP levels after stage 3 (desiccation). (A)** Schematic of the experimental plan as for fig. S12 but carried through 24 h of desiccation of droplets in air (21% $O_2$, 0.04% $CO_2$) before rehydration in various MALF formulations or 7H9. **(B)** cAMP quantification following 18 h incubation in the respective rehydration fluids. Results are mean ± SEM of three independent experiments. Results are means mean ± SEM of three independent experiments. Statistical significance was determined by two-way ANOVA followed by Tukey'smultiple comparisons test (*p < 0.05, **p < 0.01, ***p < 0.001, ****p < 0.000, ns, not significant).

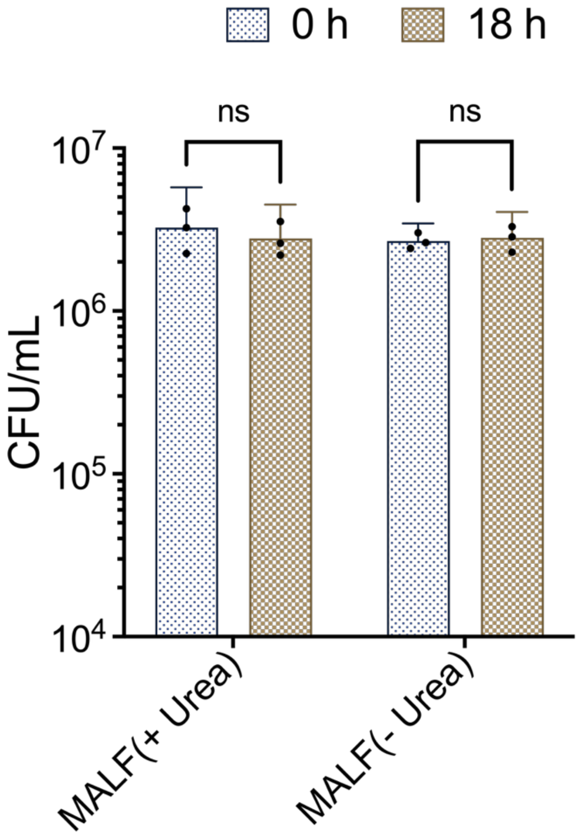


**Figure S19. Survival of desiccated Mtb in MALF with or without urea.** Viability (cfu/mL) of wild type Mtb H37Rv was measured at 0 h and 18 h post-rehydration in MALF with or without 5 mM urea. No significant change in bacterial survival was observed between time points or conditions. Results are means mean ± SEM of two independent experiments. Statistical significance was determined by two-way ANOVA followed by Tukey'smultiple comparisons test (ns, not significant).

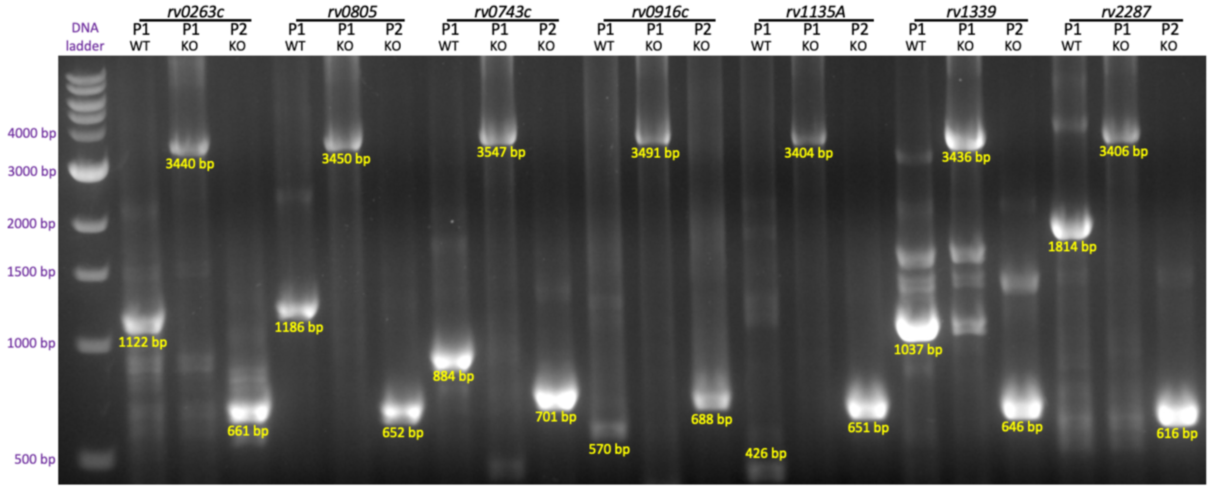
DNA ladder
rv0263c
rv0805
rv0743c
rv0916c
rv1135A
rv1339
rv2287
P1 WT
P1 KO
P2 KO
4000 bp
3000 bp
2000 bp
1500 bp
1000 bp
500 bp
3440 bp
1122 bp
661 bp
3450 bp
1186 bp
652 bp
3547 bp
884 bp
701 bp
3491 bp
570 bp
688 bp
3404 bp
426 bp
651 bp
3436 bp
1037 bp
646 bp
3406 bp
1814 bp
616 bp

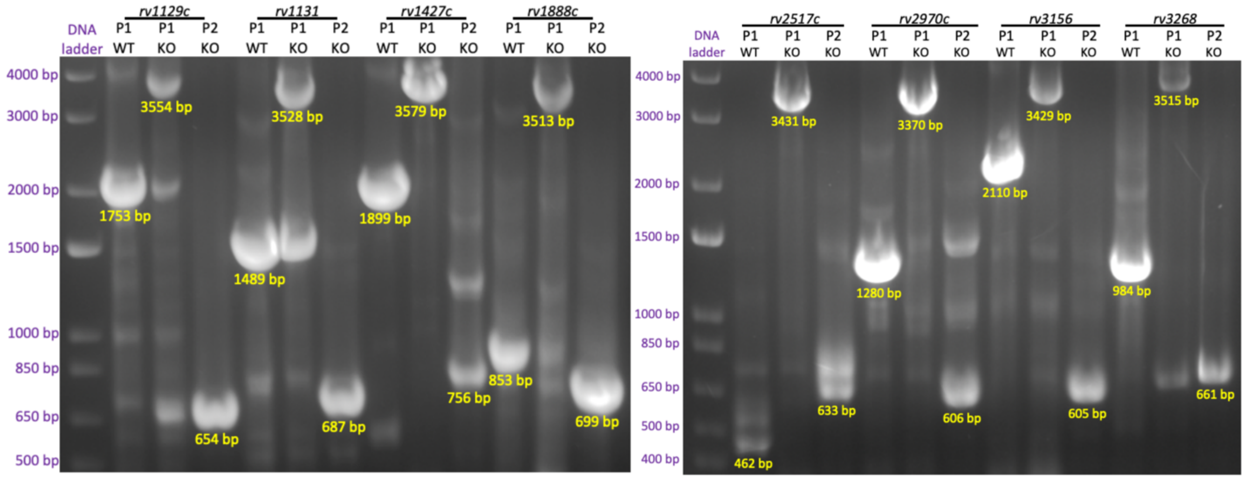
DNA ladder
rv1129c
rv1131
rv1427c
rv1888c
rv2517c
rv2970c
rv3156
rv3268
P1 WT
P1 KO
P2 KO
4000 bp
3000 bp
2000 bp
1500 bp
1000 bp
850 bp
650 bp
500 bp
400 bp
3554 bp
1753 bp
654 bp
3528 bp
1489 bp
687 bp
3579 bp
1899 bp
756 bp
3513 bp
853 bp
699 bp
3431 bp
462 bp
633 bp
3370 bp
1280 bp
606 bp
3429 bp
2110 bp
605 bp
3515 bp
984 bp
661 bp

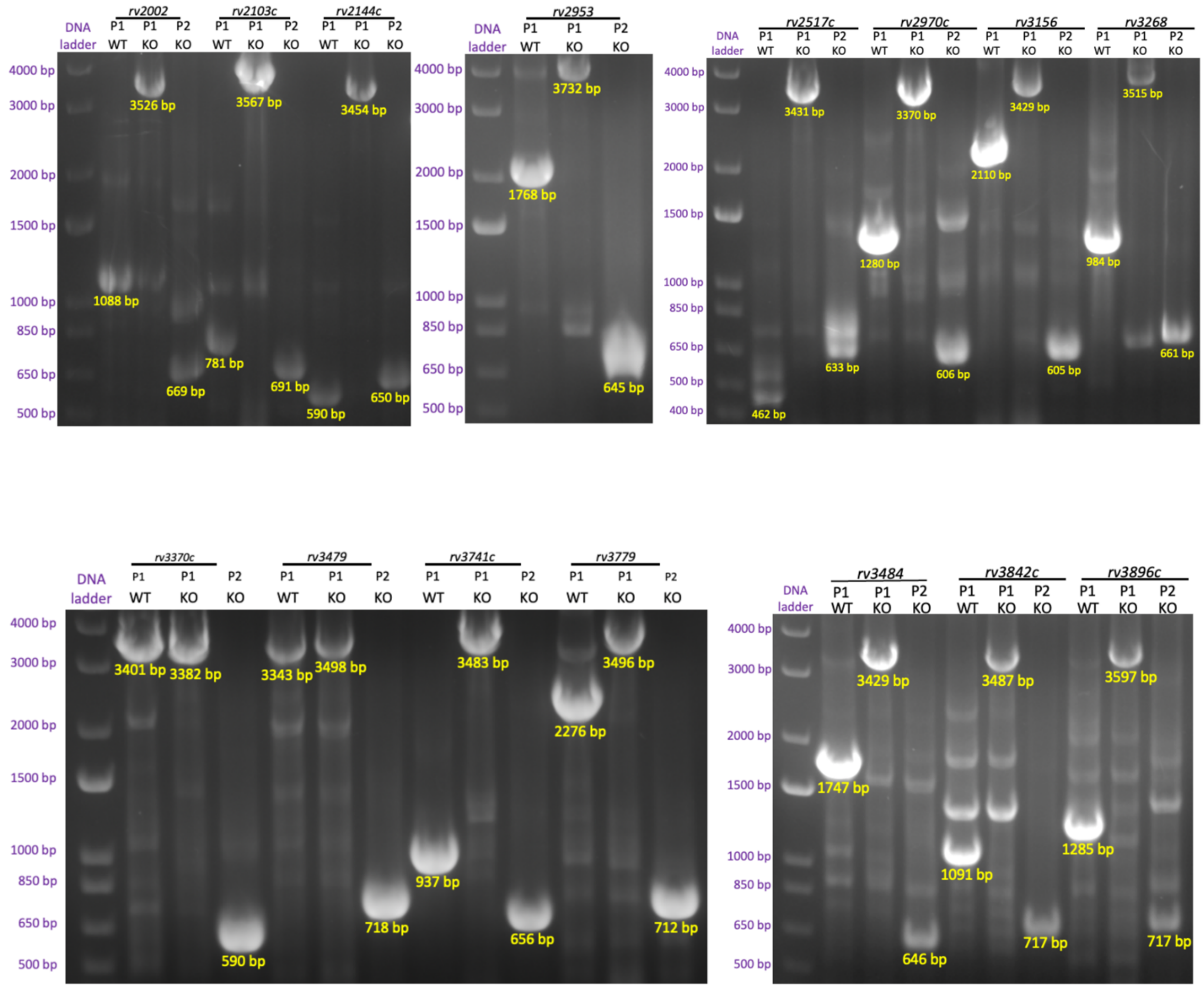


**Figure S20. PCR base confirmation of gene deletions:** Representative gel electrophoresis confirming the KO of targeted Mtb genes. For each gene, WT and KO genomic DNA were analyzed using two primer sets: Primer set 1 (P1): gene-specific forward and reverse primers. A band in the Wt lanes and its absence or size shift in the KO lanes indicates the successful loss of the native gene due to its replacement with the hygromycin resistance cassette. Primer set 2 (P2): Hygromycin-specific forward primer and gene-specific reverse primer. A band in KO lanes (yellow text) and its absence in WT lanes confirms site-specific integration of the hygromycin marker.

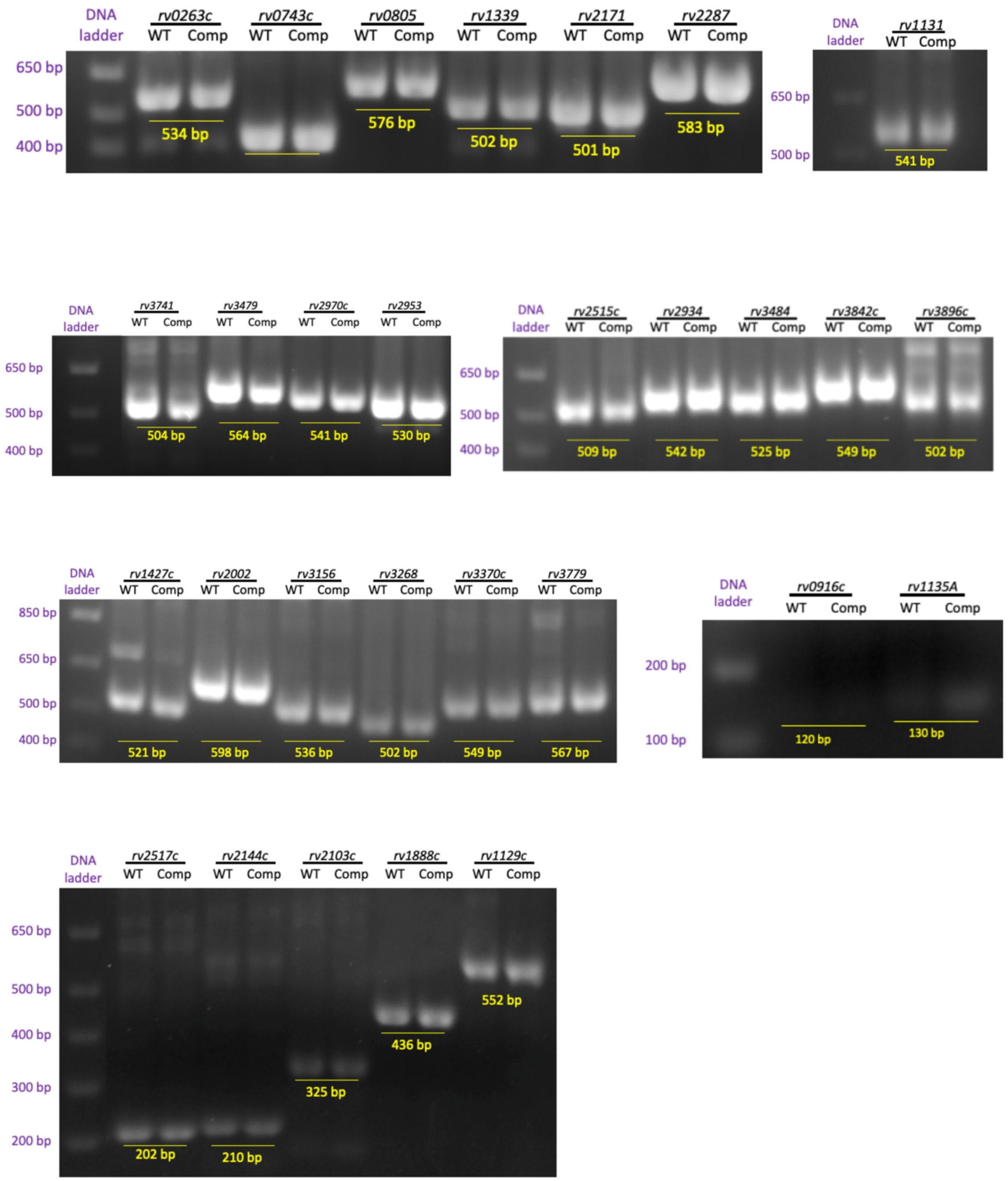


**Figure S21.** PCR Confirmation of Complemented Strains.Representative agarose gels showing the presence of targeted genes in WT and Complemented (Comp) strains. PCR was performed for each gene using gene-specific forward and reverse primers. A DNA band of the expected size (indicated in yellow) is present in both lanes, with the WT serving as a positive control. These results confirm successful gene restoration in the complemented mutants. DNA ladders (left) indicate fragment sizes.

envelope remodelling is required for Mycobacterium abscessus survival in lung infection. Nat Microbiol *8*, 481-497. 10.1038/s41564-022-01307-5.

12. Vogt, G., and Nathan, C. (2011). In vitro differentiation of human macrophages with enhanced antimycobacterial activity. J Clin Invest *121*, 3889-3901. 10.1172/JCI57235.
13. Murphy, K.C., Nelson, S.J., Nambi, S., Papavinasasundaram, K., Baer, C.E., and Sassetti, C.M. (2018). ORBIT: a New Paradigm for Genetic Engineering of Mycobacterial Chromosomes. mBio *9*. 10.1128/mBio.01467-18.
14. Beites, T., Jansen, R.S., Wang, R., Jinich, A., Rhee, K.Y., Schnappinger, D., and Ehrt, S. (2021). Multiple acyl-CoA dehydrogenase deficiency kills Mycobacterium tuberculosis in vitro and during infection. Nat Commun *12*, 6593. 10.1038/s41467-021-26941-1.
15. Nuritdinov, F., Woo, J., Schmidt, M.J., Odjourian, N.M., Cristaldo, M., Dougher, M., Antilus-Sainte, R., Heldt, T., Rhee, K., Bourouiba, L., and Gengenbacher, M. (2025). Experimental system enables studies of Mycobacterium tuberculosis during aerogenic transmission. mBio, e0095825. 10.1128/mbio.00958-25.
16. Wang, L., Wu, C., Peng, W., Zhou, Z., Zeng, J., Li, X., Yang, Y., Yu, S., Zou, Y., Huang, M., et al. (2022). A high-performance genetically encoded fluorescent indicator for in vivo cAMP imaging. Nat Commun *13*, 5363. 10.1038/s41467-022-32994-7.
17. Wilburn, K.M., Montague, C.R., Qin, B., Woods, A.K., Love, M.S., McNamara, C.W., Schultz, P.G., Southard, T.L., Huang, L., Petrassi, H.M., and VanderVen, B.C. (2022). Pharmacological and genetic activation of cAMP synthesis disrupts cholesterol utilization in Mycobacterium tuberculosis. PLoS Pathog *18*, e1009862. 10.1371/journal.ppat.1009862.
18. Wong, A.I., Beites, T., Planck, K.A., Fieweger, R.A., Eckartt, K.A., Li, S., Poulton, N.C., VanderVen, B.C., Rhee, K.Y., Schnappinger, D., et al. (2023). Cyclic AMP is a critical mediator of intrinsic drug resistance and fatty acid metabolism in M. tuberculosis. Elife *12*. 10.7554/eLife.81177.
